# Development of Paclitaxel Resistance in Triple-Negative Breast Cancer Is Associated with Extensive DNA Methylation Changes That Are Partially Reversed by Decitabine

**DOI:** 10.1101/2025.05.22.655519

**Authors:** Lenka Trnkova, Monika Burikova, Andrea Soltysova, Andrej Ficek, Jana Plava, Andrea Cumova, Lucia Rojikova, Kristina Jakic, Eva Sedlackova, Boris Tichy, Vojtech Bystry, Florence Busato, Yimin Shen, Miroslava Matuskova, Lucia Kucerova, Jorg Tost, Svetlana Miklikova, Marina Cihova, Verona Buocikova, Bozena Smolkova

**Author notes:** These authors share the last authorship (Verona Buocikova, Bozena Smolkova). Corresponding author at: Department of Molecular Oncology, Cancer Research Institute, Biomedical Research Center of the Slovak Academy of Sciences, Dubravska cesta 9, 845 05 Bratislava, Slovakia Email address (B. Smolkova).

## Abstract

**Aims:** Chemotherapy resistance remains a major challenge in breast cancer (BC) treatment. This study investigated whether resistance development is associated with DNA methylation changes and assessed the potential of the DNA methyltransferase inhibitor decitabine (DAC) to reverse these alterations and enhance chemosensitivity.

**Methods:** Molecular profiling and functional assays were used to characterize paclitaxel-(PAC) and doxorubicin-(DOX) resistant BC cell lines derived from luminal A (T-47D), triple-negative (MDA-MB-231), and HER2-positive trastuzumab-resistant (JIMT-1) models. Therapeutic responses to DAC and DOX, alone and in combination, were evaluated in MDA-MB-231 xenografts. DNA methylation–associated gene expression changes were analyzed through integrative approaches.

**Results:** Chemoresistant cells exhibited a slow-cycling phenotype, reduced tumorigenicity, and extensive genomic alterations. Upregulation of *RELB* and downregulation of *PPARG* were observed across several resistant cell lines, while *CDA* expression was uniformly elevated in all DOX-resistant models. PAC-resistant xenografts displayed widespread methylation and transcriptomic reprogramming. DAC treatment partially restored aberrant methylation patterns and increased Ki-67 expression, potentially enhancing DOX responsiveness.

**Conclusions:** Chemoresistance in BC involves extensive genomic and epigenetic reprogramming. DAC modulates methylation and tumor phenotype but is insufficient to overcome resistance, highlighting the need for rational combination strategies.

**Graphical Abstract:** 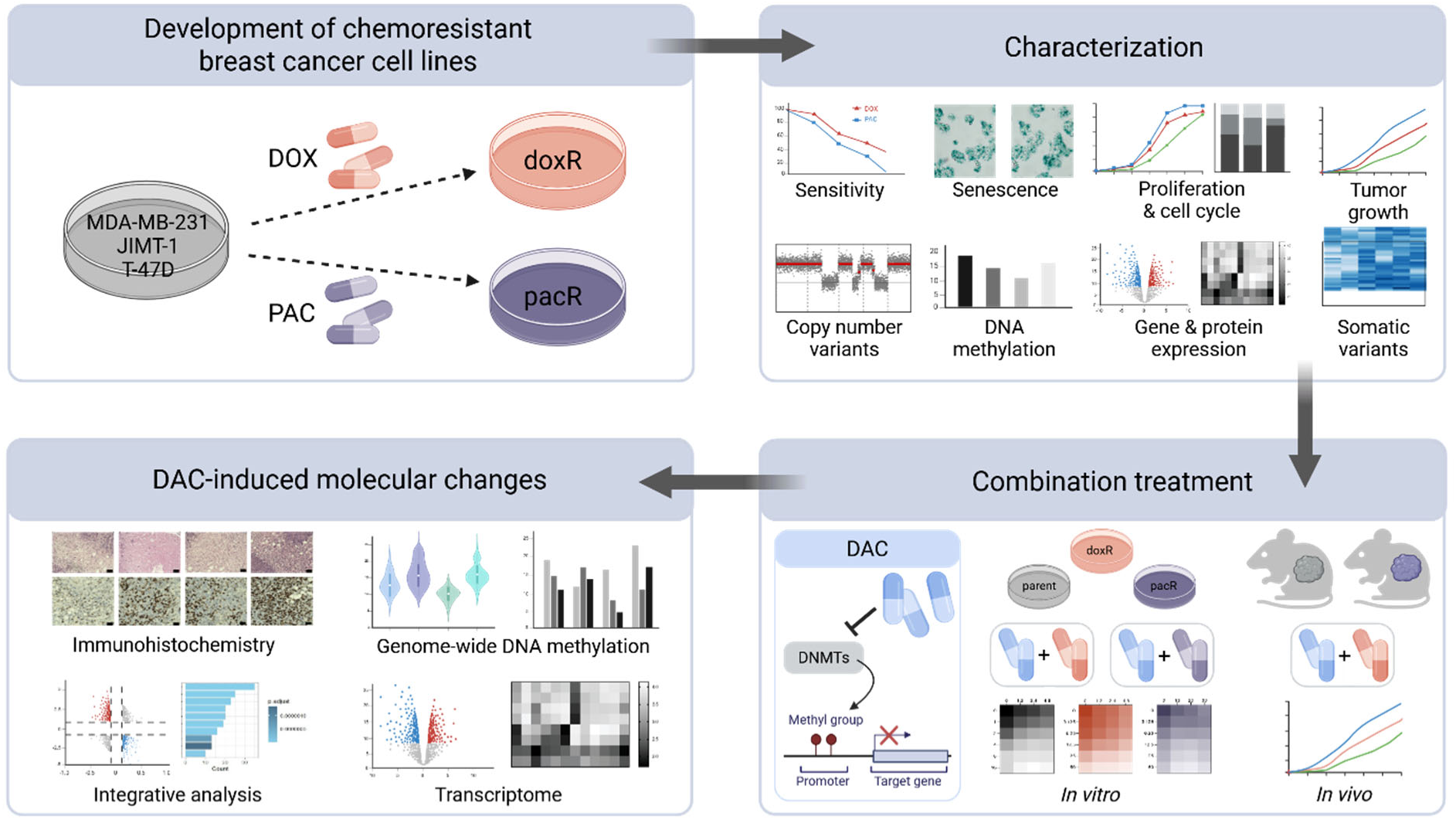

## 1. Introduction

Breast cancer (BC) remains one of the most prevalent and clinically challenging malignancies worldwide, imposing a substantial burden on public health (Bray et al., 2024). Treatment strategies are primarily guided by intrinsic surrogate subtypes, which inform therapeutic decision-making. Among these, triple-negative breast cancer (TNBC) is a highly heterogeneous and aggressive form characterized by the absence of estrogen receptor (ER), progesterone receptor (PR), and human epidermal growth factor receptor 2 (HER2) expression (Harbeck et al., 2019). The lack of these molecular targets renders TNBC particularly difficult to treat, as it is unresponsive to hormone therapies and HER2-targeting agents (Zhu et al., 2023).

Chemotherapy, the first-line treatment for TNBC, is typically reserved for later stages in other BC subtypes, such as metastatic hormone receptor-positive or HER2-positive cases (Gennari et al., 2021). Despite its widespread use, chemotherapy effectiveness is frequently hindered by the emergence of therapeutic resistance, posing a major challenge to improving patient outcomes. Resistance can emerge through various mechanisms, including alterations in drug metabolism, DNA repair pathways, and cell survival signaling (Khan et al., 2024). Increased drug efflux, primarily driven by the upregulation of ATP-binding cassette (ABC) transporters such as ABCB1 and ABCG2, plays a key role in resistance to a broad range of therapeutic agents, a phenomenon commonly referred to as multidrug resistance. Numerous recent studies have focused on targeting these transporters to overcome drug resistance (Ji et al., 2024). Additionally, alterations in key signaling pathways, including TP53, PI3K/AKT, and MAPK/ERK, as well as cellular adaptation to oxidative stress, are known to drive resistance to anthracycline-based chemotherapy, particularly doxorubicin (DOX) (Cox and Weinman, 2016; Christowitz et al., 2019; Lin et al., 2019). In contrast, resistance to microtubule-targeting agents such as paclitaxel (PAC) is often associated with changes in tubulin structure, mutations, altered tubulin expression, and posttranslational modifications (Mozzetti et al., 2005; Wattanathamsan et al., 2021; Yin et al., 2010). While several studies have described the development of PAC- or DOX-resistant BC cell lines to investigate mechanisms of chemoresistance (Paramanantham et al., 2021; Xiao et al., 2025; Zhang et al., 2024), few have involved HER2-positive and hormone receptor-positive models such as JIMT-1 and T-47D (Carr et al., 2010; Guo et al., 2025; Jeon et al., 2018).

Recent studies have underscored the importance of epigenetic modifications, particularly aberrant DNA methylation, in the development and persistence of drug-resistant phenotypes. These modifications often result in silencing tumor suppressor genes and activating oncogenes, thereby driving resistance and disease progression (Morel et al., 2020; Trnkova et al., 2024). However, genome-wide DNA methylation changes have rarely been examined in resistant BC cell lines, and to our knowledge, no relevant studies have been reported for PAC-resistant MDA-MB-231 models (He et al., 2016; Shi et al., 2020).

Targeting these epigenetic alterations with DNA methyltransferase inhibitors, such as decitabine (DAC), offers a promising approach to restoring chemosensitivity in solid tumors (Hu et al., 2021; Jin et al., 2021; Morel et al., 2020). Due to the challenges of high-dose DAC regimens, several clinical trials are investigating the combination of low-dose DAC with other therapeutic approaches to evaluate efficacy and safety (NCT03875287, NCT02961101, NCT01799083, NCT05638984, NCT04611711, NCT05320640). While some trials have reported promising outcomes, others are ongoing, and comprehensive results are pending. Further research is essential to fully understand the benefits and limitations of these strategies.

Previously, we showed that DAC enhances DOX sensitivity in JIMT-1 and T-47D BC cells, affecting DNA methylation and gene expression. In MDA-MB-231 spheroids, the DAC and DOX combination synergistically reduced cell migration (Buocikova et al., 2022a). Building on these results, we extended our investigation to chemotherapy-resistant BC cell lines, focusing on DNA methylation changes associated with the development of resistance. The primary goal of this study was to evaluate DAC’s capacity to normalize aberrant DNA methylation patterns in drug-resistant BC tumors. To generate resistant derivatives, we employed two chemotherapeutic agents commonly used in the treatment of TNBC and other aggressive BC subtypes, DOX and PAC. Derived cell lines were characterized at both cellular and molecular levels and used to establish xenograft models for evaluating their tumorigenicity and phenotypic features. Finally, using a multi-omics approach, we assessed DAC’s ability to normalize DNA methylation profiles associated with the development of PAC resistance in the TNBC model *in vivo*.

## 2. Materials and Methods

### 2.1. Cell culture and development of chemotherapy-resistant cells

Cell culture experiments were conducted using three human BC cell lines of distinct subtypes: MDA-MB-231 (HTB-26™, ATCC), JIMT-1 (ACC 589, German Collection of Microorganisms and Cell Cultures GmbH), and T-47D (HTB-133™, ATCC). The cells were cultured in Dulbeccós Modified Eaglés Medium (DMEM) high glucose (Sigma-Aldrich, Germany). The cell culture medium was supplemented with 10% fetal bovine serum (Sigma-Aldrich, Germany), 2 mM glutamine (PAA Laboratories GmbH, Germany), HyClone penicillin-streptomycin solution (Cytiva Austria GmbH, Austria), and 2.5 μg/mL amphotericin B (Sigma-Aldrich, Germany).

DOX-resistant (doxR) and PAC-resistant (pacR) cells were derived from MDA-MB-231, JIMT-1, and T-47D cell lines through continuous exposure to DOX (Sigma-Aldrich, Germany) or PAC (Selleck Chemicals, Houston, USA), respectively. DoxR cells were exposed to an initial concentration of 1 ng/mL DOX for 24 h, after which the cell culture medium was changed to allow for cell recovery and growth. Upon reaching confluence, cells were passaged, and the DOX concentration was increased by 1 ng/mL the following day. This process was repeated until reaching the final DOX concentration of 10 ng/mL for MDA-MB-231, 31 ng/mL for JIMT-1, and 17 ng/mL for T-47D cells.

PacR cells were exposed to an initial concentration of 1 ng/mL PAC for several days until reaching confluence. Following passaging, cells were allowed to recover in fresh drug-free medium. The next day, PAC concentration was increased by 1 ng/mL. This process continued until the final PAC concentrations of 20 ng/mL for MDA-MB-231 and 19 ng/mL for JIMT-1 cells were reached. Unexposed parental cells were cultured in parallel during the derivation process. Attempts to derive the T-47D pacR cell line were unsuccessful.

After attaining the final drug concentrations, corresponding to approximately a 5-fold decrease in cell drug sensitivity, the cells were maintained for a minimum of two months without further exposure to these agents. This period ensured that any changes observed in subsequent experiments reflected stable adaptations acquired by the resistant cells. Short tandem repeat (STR) profiling was performed following this period to verify cell identity.

### 2.2. Cell viability assay

The sensitivity of parental and chemotherapy-resistant cells to selected drugs was determined by measurement of cell viability upon drug exposure. Cells were seeded in a 96-well cell culture plate (Sarstedt, Germany) and treated for six days with varying concentrations of PAC (1 - 16 ng/mL), DOX (6.25 - 100 ng/mL) or DAC (0.2 - 500 ng/mL), covering the viability range of the parental cells. MDA-MB-231 parental and chemotherapy-resistant cells were also exposed to combination treatment as follows: after 24 h of DAC treatment in concentrations corresponding to IC_20_ (1.2 - 4.8 ng/mL for MDA-MB-231 parent and MDA-MB-231 doxR; 10 - 30 ng/mL for MDA-MB-231 pacR), cells were exposed to PAC (1 - 16 ng/mL) or DOX (3.125 - 50 ng/mL) for 5 days. Cell viability was measured using CellTiter-Glo® Luminescent Cell Viability Assay (Promega Corporation, Wisconsin, USA), and cell luminescence was detected on a GloMax® Discover Microplate Reader (Promega Corporation, Wisconsin, USA). Cell viability was determined as a percentage of cell luminescence relative to control (unexposed cells, set to 100%). Data are represented as mean ± standard deviation (SD) from three independent experiments performed in triplicates. CalcuSyn software (Biosoft, UK) was used to compute the IC_50_ values and combination index (CI) (Chou, 2006). CI represents the effect of combined therapy: synergism (CI < 1), additivity (CI = 1), or antagonism (CI > 1).

### 2.3. Cell proliferation

Cells were seeded in a 96-well plate (TPP, Switzerland) at the following concentrations: 1.5 × 10^3^ cells/well for MDA-MB-231 parental, doxR, and pacR, 3 × 10^3^ cells/well for JIMT-1 parental, doxR, and pacR, 3.7 × 10^3^ cells/well for T-47D parental and doxR. Cell proliferation was determined by analysis of cell confluence measured for 7 days using IncuCyte ZOOM™ Live-Cell Imaging System (Essen BioScience, UK) and IncuCyte ZOOM 2016A software. Results are presented as mean ± SD from three independent experiments performed in hexaplicates.

### 2.4. Doubling time

Cells were seeded in 6-well plates (TPP, Switzerland) at the following concentrations: 4 × 10^4^ cells/well MDA-MB-231 parent, doxR and pacR, 8.5 × 10^4^ cells/well JIMT-1 parent, doxR and pacR, 1.7 × 10^5^ cells/well T-47D parent and doxR, and cultured for 96 h. Afterward, the number of cells was counted using CellDropTM (DeNovix, Delaware, USA). The cell doubling time was calculated using the formula DT=t/(ln(N_t_/N_i_)ln2), where DT is doubling time in hours (h), t is culture time (h), N_t_ is the cell number at t, and N_i_ is the initial cell number (Rainaldi et al., 1991). Results are presented as mean ± SD from three independent experiments.

### 2.5. Cell cycle analysis

2×10^5^ cells were resuspended in 0.05% Triton X-100 (Sigma-Aldrich, Germany) and treated with 10 µg/mL Ribonuclease A (Sigma-Aldrich, Germany) for 20 min at 37°C. Subsequently, the cells were incubated at 4°C for 1 min, and nucleic acids were stained with 40 µg/mL propidium iodide (Sigma Aldrich, Germany). Cell cycle analysis was performed using a Cytoflex S flow cytometer (Beckman Coulter, California, USA). Data were analyzed using FCS Express (De Novo Software, California, USA).

### 2.6. Cell senescence

β-galactosidase activity in both parental and chemotherapy-resistant cells was assessed using the Senescence β-Galactosidase Staining Kit (Cell Signaling Technology, Massachusetts, USA). This assay detects β-galactosidase, an enzyme expressed in senescent cells, which becomes active at pH 6. The enzyme cleaves the substrate 5-bromo-4-chloro-3-indoyl β-D-galactopyranoside, resulting in the development of a visually detectable blue color. Microscopic images of stained cells were captured using the Axio Vert.A1 ZEISS light microscope and processed with Zen 2.6 software (Carl Zeiss Microscopy GmbH, Germany). β-galactosidase staining was independently evaluated by two observers (AC and MB), and staining intensity was assigned using a semi-quantitative scoring system: + (score 1), ++ (score 2), and +++ (score 3).

### 2.7. Cell migration

MDA-MB-231 parent and doxR (3×10^4^ cells/well), pacR (3.5×10^4^ cells/well) cells, JIMT-1 parent and doxR (3×10^4^ cells/well), pacR (5×10^4^ cells/well) cells, and T-47D parent and doxR (6×10^4^ cells/well) cells were seeded into Incucyte® Imagelock 96-well Plate (Sartorius, Germany). The scratch wound was performed using the Incucyte® 96-Well Woundmaker Tool (Sartorius, Germany) after 24 h, according to the manufacturer’s instructions. Cells were maintained in a culture medium. Analysis of wound confluence was performed using IncuCyte ZOOM™ Live-Cell Imaging System (Essen BioScience, UK) and IncuCyte ZOOM 2016A software. Data are presented as mean ± SD from three independent experiments.

### 2.8. High-resolution array comparative genomic hybridization

DNA was isolated from cell pellets using a NucleoSpin® Tissue kit (Macherey-Nagel, Germany) according to the manufacturer’s instructions. DNA concentration and quality were determined using a NanoDrop 1000 spectrophotometer (Thermo Fisher Scientific, USA). High-resolution array comparative genomic hybridization combined with single nucleotide polymorphism (aCGH+SNP) analysis was conducted using the Agilent platform (Agilent Technologies, USA). In brief, 750 ng of DNA was digested and labeled using the SureTag Complete DNA Labeling Kit (Agilent Technologies, California, USA) in accordance with the manufacturer’s protocol. Labeled DNA, along with the Female Human Reference DNA provided in the kit, was combined and loaded on Agilent SurePrint G3 Human Genome CGH+SNP Microarray Kit, 2×400K, and hybridized for 40 hours at 67°C. Post-hybridization, slides were washed and scanned using a SureScan Microarray Scanner (Agilent Technologies, California, USA). Data were further processed using the Bioconductor rCGH package version 1.36.0 (Commo, 2024). Signals were adjusted for GC content and Cy3/Cy5 bias, followed by log_2_(relative ratio) (LRR) computation. A Circular Binary Segmentation algorithm and an expectation-maximization algorithm were applied to segment genome profiles and for LRR centering, respectively. UCSC hg19 RefSeq annotations and genome locations were used to map genes to the segmented genome profiles using the Bioconductor GenomicRanges package version 1.58.0 (Lawrence et al., 2013). Differentially altered genomic segments between parental and resistant cells were identified based on an absolute LRR value difference greater than 0.5.

### 2.9. Whole-exome sequencing

DNA from cell pellets was extracted as described above. A total of 500 ng of input DNA was utilized for library preparation with the xGen™ DNA Lib Prep EZ UNI kit (IDT, Belgium), with library quantification performed via the KAPA Library Quantification Kit (Roche, Switzerland). The pooled libraries were sequenced in paired-end 2×150 mode on a DNBSEQ-G400 sequencer (MGI, Germany). The Burrows-Wheeler Aligner (Li and Durbin, 2009) was employed to align preprocessed reads to the reference human genome (GRCh38-p10). Somatic variants were identified in parental and corresponding therapy-resistant cell lines by the SomaticSeq pipeline (Fang, 2020). To identify prevalent variants in chemotherapy-resistant cell lines, the data were filtered for tumor variants with a frequency exceeding 0.35. Acquired potentially deleterious variants were selected based on tumor variant frequency above 0.35 and a combined annotation-dependent depletion phred-like (CADD PHRED) score greater than 20. The data were summarized using the oncoplot created with the R package *maftools* version 2.22.0 (Mayakonda et al., 2018).

### 2.10. Human Cancer Drug Resistance RT² Profiler PCR Array

To investigate drug resistance mechanisms, the Human Cancer Drug Resistance RT² Profiler PCR Array (PAHS-004ZD, Qiagen, Germany) was employed for the expression analysis of 84 resistance-related genes. RNA was extracted from cell pellets with the RNeasy® Mini Kit (Qiagen, Germany), and on-column DNA digestion was performed with the RNase-Free DNase Set (Qiagen, Germany). The RNA concentration and quality were assessed on a NanoDrop 1000 spectrophotometer (Thermo Fisher Scientific, USA). For cDNA synthesis, 500 ng of purified RNA was processed with the RT² First Strand Kit (Qiagen, Germany). The resulting cDNA served as the template for reverse transcription PCR (RT-PCR), conducted with RT² SYBR® Green Mastermix (Qiagen, Germany) and the Human Cancer Drug Resistance RT² Profiler PCR Array. RT-PCR experiments were carried out on a Bio-Rad CFX96™ Real-Time PCR System (Bio-Rad, California, USA). Data from three independent experiments were processed and analyzed via the online GeneGlobe Data Analysis Center (Qiagen, Germany).

### 2.11. RT-qPCR analysis of key DNA methylation regulators and DAC metabolism-related genes

Total RNA (2000 ng) was used for cDNA synthesis using the RevertAid H Minus First Strand cDNA Synthesis Kit (Thermo Fisher Scientific, USA). Expression of selected genes was quantified by quantitative reverse transcription PCR (RT-qPCR) using TaqMan gene expression assays: *DNMT1* (Hs00945875_m1), *DNMT3A* (Hs01027166_m1), *DNMT3B* (Hs00171876_m1), *TET1* (Hs00286756_m1), *TET2* (Hs00325999_m1), *TET3* (Hs00896441_m1), *CDA* (Hs00156401_m1) and *GAPDH* (Hs99999905_m1). RT-qPCR was performed on a Bio-Rad CFX96™ real-time cycler using TaqMan Gene Expression Master Mix (Applied Biosystems, USA) and 50 ng of cDNA input, following the manufacturer’s instructions. Individually designed primers were used to analyze *DCK* expression (5′-TCTGAGGGGACCCGCATCAA-3′ and 5′-TGCACCATCTGGCAACAGGTT-3’) and *RPLP0* (5′-AGGAAACTCTGCATTCTCGCT -3′ and 5′-CAAGAAGGCCTTGACCTTTTCA-3’). 50 ng of cDNA and GoTaq qPCR MasterMix (Promega, USA) were used for analysis. Relative quantification of gene expression was determined using the ΔCT method, with *GAPDH* or *RPLP0* as the reference gene, and fold change (FC) was calculated using the ΔΔCT method. Data are presented as mean ± SD from three independent experiments, each performed in triplicate.

### 2.12. Western blot

Proteins were isolated from cell pellets of parental and chemotherapy-resistant cells using RIPA buffer (Cell Signaling Technology, USA) supplemented with PhosStop (Roche, Switzerland) and cOmplete Protease Inhibitor Cocktail (Roche, Switzerland). Protein concentration was determined using the Pierce BCA protein assay (Thermo Fisher Scientific, USA). Thirty µg of proteins were separated using SDS-PAGE gel electrophoresis and transferred to a nitrocellulose membrane (Bio-Rad, Germany). Membranes were subsequently blocked for 1 h in 5% non-fat milk (PanReac Applichem, Germany) diluted in Tris buffered saline (TBS). Membranes were incubated with CDA (cat. no. PA5-84630, Invitrogen, USA), DHFR (cat. no. 43497, Cell Signaling Technology, USA) or DCK (cat. no. PA5-27787, Invitrogen, USA) primary antibodies at 4°C overnight and Goat anti-Rabbit IgG (H+L) Highly Cross-Adsorbed Secondary Antibody, Alexa Fluor™ 680 (Invitrogen, USA) for 1 h at room temperature (RT). Protein stains were detected using the Odyssey® Fc imaging system (LI-COR Biotechnology, USA) at 700 nm. Membranes were subsequently incubated with GAPDH (cat. no. 97166, Cell Signaling Technology, USA) or β-actin (cat. no. A1978, Sigma-Aldrich, Germany) primary antibodies for 1 h at RT followed by incubation with anti-mouse IgG, HRP-linked secondary antibody (Cell Signaling Technology, USA) for 1 h at RT. Washing steps with TBS supplemented with 0.1 % Tween (Sigma-Aldrich, Missouri, USA) (TBST) were applied between the antibody incubations. Finally, membranes were incubated for 5 min with SuperSignal West Dura Extended Duration Substrate (Thermo Fisher Scientific, IL, USA), and chemiluminescent protein staining was detected using Odyssey® Fc imaging system (LI-COR Biotechnology, USA). Protein quantification was performed via ImageJ software. Relative protein levels of chemotherapy-resistant cells were normalized towards their parental counterpart (protein level set to 1). CDA protein levels in T-47D cells were not detected by the selected primary antibody (Buocikova et al., 2022b). Data represent mean ± SD from three independent experiments.

### 2.13. Global DNA methylation

Global DNA methylation levels were assessed by quantifying the methylation of LINE-1 retrotransposable elements in the DNA of the studied cell lines by pyrosequencing using PyroMark® Q24 CpG LINE-1 Kit (Qiagen, Germany). DNA was extracted with the FlexiGene® DNA Kit (Qiagen, Germany). Bisulfite modification of 2000 ng of extracted DNA was performed using the EpiTect Bisulfite Kit (Qiagen, Germany). The LINE-1 sequence was amplified by PCR with PyroMark® Q24 CpG LINE-1 and the PyroMark® PCR Kit (Qiagen, Germany). Pyrosequencing was conducted using PyroMark® Gold Q24 Reagents and the PyroMark® Q24 system (Qiagen, Germany). Data were analyzed with PyroMark Q24 2.0.6 software (Qiagen, Germany). The methylation levels of three CpG sites within the LINE-1 sequence were averaged, and results are presented as mean ± SD from three independent experiments.

### 2.14. Tumorigenicity in vivo

*In vivo* experiments were conducted using female SCID beige mice (SCID/bg; Charles River, Germany). The mice were housed in the authorized animal facility (SK UCH 02022). The Institutional Ethical Committee and the State Veterinary and Food Administration of the Slovak Republic approved the study under registration number 3616/18-221/3. All animal experiments were carried out in accordance with institutional guidelines and approved protocols.

For tumorigenicity testing, 1.5 × 10^6^ MDA-MB-231 parent, doxR, pacR and JIMT-1 parent, doxR, and pacR cells were resuspended in 50 µL DMEM high glucose diluted with ECM Gel from Engelbreth-Holm-Swarm murine sarcoma (Sigma-Aldrich, Missouri, USA) (1:1) and bilaterally orthotopically injected into the fourth mammary fat pad of SCID/bg mice. T-47D cells were not included in the *in vivo* experiments due to their inherently low tumorigenic potential, requiring substantially higher cell numbers for tumor establishment and dependence on exogenous estrogen supplementation (Shishido and Nguyen, 2012). Tumor length and width were regularly measured by caliper, and tumor volumes were calculated using the formula V = 1/6 π × L × W x (L + W)/2, where V is tumor volume (mm^3^), L is tumor length (mm), and W is tumor width (mm) (Rodallec et al., 2022). Mice were sacrificed at the humane endpoint, defined as the time when at least one of the tumors in the group reached a size of 1000 mm^3^, ulceration of the tumor, significant weight loss exceeding 15% of initial body weight, signs of distress such as lethargy, impaired mobility, or inability to access food and water appeared. Animals were monitored daily, and decisions were made in accordance with the institutional guidelines and ethical approvals. The weight of the tumors was determined upon autopsy.

### 2.15. Combination treatment in vivo

For combination treatment, orthotopic xenografts were induced using 2.5 × 10^5^ MDA-MB-231 parental and 1.5 × 10^6^ MDA-MB-231 pacR cells resuspended in 50 µL DMEM high glucose diluted with ECM Gel from Engelbreth-Holm-Swarm murine sarcoma (Sigma-Aldrich, Missouri, USA) (1:1). Cells were injected bilaterally into the fourth mammary fat pad of SCID/bg immunodeficient mice. Mice were randomly distributed into four treatment groups: vehicle, DAC, DOX, and the combination treatment. The treatment started upon the development of palpable tumors (day 6 for mice injected with MDA-MB-231 parental cells and day 16 for mice injected with MDA-MB-231 pacR cells). The 26-day treatment consisted of two 12-day cycles separated by a 2-day drug-free interval (Fig. S1A). Each cycle involved intraperitoneal treatment with either the vehicle consisting of 0.16% dimethyl sulfoxide (DMSO; Sigma-Aldrich, Missouri, USA) or saline (Fresenius Kabi, Czech Republic), 0.4 mg/kg DAC (MedChem Express, China), 1 mg/kg DOX (medac GmbH, Germany), or a combination of DAC and DOX. Tumor growth was regularly measured by caliper, and volumes were calculated according to the formula described above. Mice were sacrificed one day after completing the two treatment cycles, and tumor weight was determined upon autopsy. Tumor specimens were then collected for downstream molecular analyses. The prolonged mono- and combination therapy with DAC and DOX in MDA-MB-231 pacR xenografts started on day 6 after the cancer cell injection and followed a 39-day treatment schedule (Fig. S1B). The treatment scheme for the pilot experiment in MDA-MB-231 doxR xenografts is illustrated in Fig. S1C.

### 2.16. Histology & immunohistochemistry

Tumor specimens were fixed in 10% neutral buffered formalin solution (Sigma-Aldrich, Missouri, USA) at RT for 24 hours, embedded in paraffin, and sectioned at 4 µm thickness. Sections were stained with hematoxylin and eosin (H&E, Sigma-Aldrich, Missouri, USA), and the extent of necrosis was evaluated by the pathologist under light microscopy (Mateo FL, Leica, Germany). For immunohistochemical (IHC) analysis, deparaffinization, rehydration, and epitope retrieval using high-pH Target Retrieval Solution (DAKO, Denmark) were carried out in a PT Link module (DAKO, Denmark) at 96°C for 20 min. Slides were then washed with FLEX Wash Buffer (DAKO, Denmark) and processed using the automated DAKO Autostainer Link 48. Endogenous peroxidase activity was blocked by incubating the sections with FLEX Peroxidase Block (DAKO, Denmark) for 5 min. Tissue sections were incubated at RT for 20 min with anti-human primary antibodies against Ki67 (clone MIB-1), Estrogen Receptor α (ERα, clone EP1), and Progesterone Receptor (clone PgR 636) (FLEX, DAKO, Denmark). For CD31 detection, tumor sections were incubated overnight at 4°C with an anti-CD31 antibody (clone EPR17259, Abcam, UK) at a 1:2000 dilution. This was followed by a 15-minute incubation with the DakoCytomation EnVision+ System-HRP. Positive staining was visualized using 3,3’-Diaminobenzidine (DAB substrate-chromogen solution, DAKO, Denmark) for 5 min. Counterstaining was performed with hematoxylin (FLEX, DAKO, Denmark) for 5–10 min. Slides were washed with FLEX Wash Buffer (DAKO, Denmark) between steps and mounted using Faramount Mounting Medium (DAKO, Denmark). The staining evaluation was conducted under light microscopy (Mateo FL microscope, Leica, Germany). For HER2 staining, the HercepTest kit (DAKO, Denmark) was employed on the DAKO Autostained Link platform, following a similar protocol except for tissue dehydration. Epitope retrieval was performed using the PT Link module (DAKO, Denmark) at 97°C for 40 min.

### 2.17. RNA Sequencing

Snap-frozen tumor tissue (up to 30 mg) was mechanically homogenized in liquid nitrogen, and RNA from tumor tissue was isolated using RNeasy® Mini Kit (Qiagen, Germany). On-column DNA digestion was performed using the RNase-Free DNase Set (Qiagen, Germany). RNA concentration and quality were assessed by NanoDrop spectrophotometer (Thermo Fisher Scientific, USA).

The library was prepared using 500 ng of total RNA as an input and QuantSeq 3’ mRNA-Seq V2 Library Prep Kit with UDI (Lexogen, Austria) with UMI Second Strand Synthesis Module for QuantSeq FWD (Lexogen, Austria). Libraries were sequenced in single-end mode using Illumina technology. The sequencing depth varied between 11 – 13 million reads per sample. Quality-checked (FastQC, MultiQC, minion, swan) and preprocessed reads (Trimmomatic, FastQC, MultiQC) were mapped (STAR) to the reference human genome with gene annotation (GRCh38-p10). FeatureCounts was used to count and summarize mapped reads to genes. Differential gene expression was performed using DESeq2 (Love et al., 2014).

Cut-off values for differentially expressed genes (DEGs) were set to |FC| > 1.5 and adjusted p-value < 0.05. Pathway analysis was performed using g:Profiler web server (Kolberg et al., 2023) (accessed May 16, 2025).

### 2.18. Genome-wide DNA methylation analysis in tumor xenografts

Tumor tissue (up to 20 mg) was manually homogenized in liquid nitrogen, and DNA was isolated using a NucleoSpin® Tissue kit (Macherey-Nagel, Germany). DNA concentration was determined using a NanoDrop 1000 spectrophotometer (Thermo Fisher Scientific, USA). 1 µg of genomic DNA was bisulfite-treated with the EpiTect® Fast DNA Bisulfite Kit (Qiagen, Germany). Genome-wide DNA methylation was analyzed on > 850 000 CpGs with the Infinium Human Methylation EPIC v1.0 BeadChip (Illumina, San Diego, USA). Data analysis was performed using the Bioconductor ChAMP package version 2.22 (Morris et al., 2014; Tian et al., 2017). All samples passed quality control (less than 10% of probes with a detection p-value >0.01). Probes with a detection p-value > 0.01 in at least 2% of the samples, with a beadcount <3 in at least 5% of samples, non-CpG probes and multi-hit probes (as described in Nordlund et al., 2013) were filtered. A total of 18533 probes matching the mouse genome (Mus musculus GRCm39; complete match and up to 3 mismatches) were removed using a previously reported approach (Pichon et al., 2021). β values representing methylation proportions, defined as a ratio of intensities between methylated and combined methylated and unmethylated bead types and ranging from 0 – 1 were normalized using Beta MIxture Quantile (BMIQ) normalization (Teschendorff et al., 2013). Differentially methylated CPGs (DMPs) with an adjusted p-value < 0.05 between DAC- and vehicle-treated MDA-MB-231 parental and pacR xenografts (Benjamini-Hochberg method) were determined with champ.DMP function. The proportion of DMPs was calculated by normalizing the percentage of DMPs across genomic features and CpG islands to the percentage of corresponding probes in the Human Infinium Methylation EPIC v1.0 BeadChip array.

### 2.19. Integrative analysis

An integrative analysis of transcriptomic and methylomic data was conducted to investigate the impact of DNA methylation on gene expression. Pearson correlation coefficients (r) were calculated between the β values of DMPs (with |Δβ| > 0.1) located in functional genomic regions (excluding open sea regions) and the normalized expression levels of the corresponding genes with |FC| >= 1.5. A significant correlation was defined as an inverse correlation coefficient of r < -0.3 and an adjusted p-value < 0.05, based on established criteria (Landau and Everitt, 2003; Xu et al., 2019).

### 2.20. Statistical analysis

A Shapiro-Wilk test was used to test the normal distribution of the data. Statistical analysis of cell viability, cell cycle, and comparison of tumor weights was performed using one-way ANOVA followed by relevant multiple comparison tests or Kruskal-Wallis test with a Dunn multiple comparison test in the case of non-normally distributed data. An unpaired t-test was used for statistical analysis of doubling time, RT-qPCR, and Western blot. Mann-Whitney U test was used for statistical analysis of β-galactosidase staining. A general linear model for repeated measures, incorporating Greenhouse-Geisser correction when assumptions of sphericity were violated, was used to assess the differences in cell proliferation, cell migration, and the impacts of four treatment types across various time points. Statistical analysis was performed using GraphPad Prism 10.4.1 and SPSS software package version 23 (IBM SPSS, Inc., Chicago, IL, US).

## 3. Results

### 3.1. Chemotherapy-resistant cells exhibit distinct morphology and slow-cycling phenotype

To generate chemotherapy-resistant cell lines, MDA-MB-231, JIMT-1, and T-47D cells were exposed to incremental concentrations of DOX and PAC over a period of several months until reaching the final DOX/PAC concentrations described in Fig. S2A. This process allowed for the selection of subpopulations with enhanced survival capabilities under drug pressure. Phase-contrast microscopy revealed distinct morphological changes in the resistant cell lines compared to their parental counterparts (Fig. S2B). MDA-MB-231 doxR cells appeared more compact than the MDA-MB-231 parental cells, while MDA-MB-231 pacR cells tended to form aggregates upon passage. JIMT-1 pacR cells exhibited an elongated, mesenchymal-like morphology. Cell viability following treatment with DOX and PAC confirmed the successful development of resistant cell lines (Fig. 1A). IC_50_ FC values for DOX were 5.3 in MDA-MB-231 doxR, 5.6 in JIMT-1 doxR, and 7.8 in T-47D doxR cells, while for PAC, IC_50_ FC values were 4.7 in MDA-MB-231 pacR and 6.3 in JIMT-1 pacR cells, all relative to their respective parental cell lines.

**Fig. 1.**
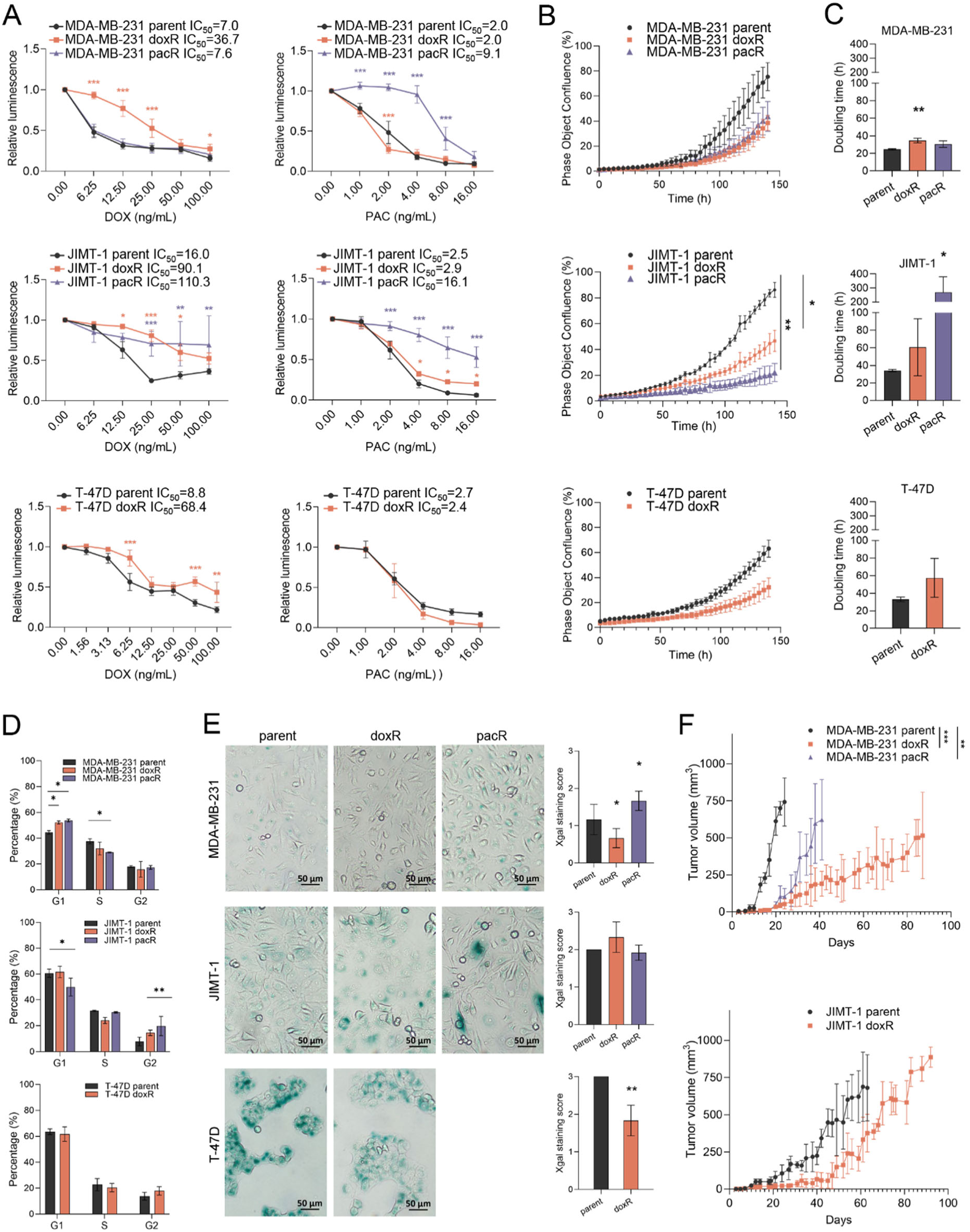
Chemoresistant cells show reduced proliferative capacity, an extended doubling time *in vitro*, and slower tumor growth *in vivo*. (A) Cell viability values of parental and resistant cell lines after 6 days of DOX or PAC treatment expressed relative to the non-treated control (set to 1). Statistical significance was assessed relative to treated parental cell lines. Data represent mean ± SD from three independent experiments. IC_50_ values were calculated using the CalsuSyn software. B) The proliferation of parental and chemotherapy-resistant cells assessed by the IncuCyte^TM^ ZOOM system presented as cell confluence (%) over time. (C) Doubling time of parental and resistant cells. Data represent the mean ± SD from three independent experiments, with statistical significance assessed relative to parental cells. (D) Cell cycle distribution of parental and resistant cells, determined by flow cytometry, with data shown as mean ± SD from three independent experiments. (E) β-galactosidase staining of parental and resistant cells. Representative images from three independent experiments at 200× magnification along with evaluation of staining intensity using a semi-quantitative scoring system; data represent mean ± SD from three independent experiments. (F) *In vivo* tumorigenicity of parental and resistant cell lines. Tumor growth following orthotopic injection of MDA-MB-231 and JIMT-1 cells into SCID/bg mice. Data represent mean ± SD from six tumors per group. * p < 0.05, ** p < 0.01, *** p < 0.001

Proliferation analysis using the IncuCyte^TM^ ZOOM system revealed a significant reduction in growth rates for JIMT-1 doxR and pacR cells, consistent with a slow-cycling phenotype (Fig. 1B). This was further supported by prolonged doubling times in MDA-MB-231 doxR and JIMT-1 pacR cells (Fig. 1C). Despite these changes in proliferation, cell migration remained unaffected in resistant derivatives (Fig. S2C, D), suggesting that resistance-associated adaptations predominantly influence cell cycle dynamics rather than motility. Flow cytometry further confirmed this phenotype, showing G1 phase accumulation in MDA-MB-231 doxR and pacR cells and a G2 phase shift in JIMT-1 pacR cells (Fig. 1D, Fig. S2E). In contrast, T-47D doxR cells displayed a cell cycle distribution comparable to the parental cells. Additional insight was gained from β-galactosidase staining, a marker of senescence. While MDA-MB-231 doxR and T-47D doxR cells exhibited decreased β-galactosidase activity, MDA-MB-231 pacR cells showed elevated levels (Fig. 1E). Consistent with these findings, resistant cells exhibited markedly reduced tumorigenic potential *in vivo*, as reflected by delayed tumor growth and extended time to humane endpoint (Fig. 1F). MDA-MB-231 parental cells exhibited the highest tumorigenicity, reaching the endpoint by day 24, while their doxR and pacR derivatives showed substantially slower progression (days 87 and 41, respectively). JIMT-1 doxR tumors followed a similar pattern, reaching the endpoint at day 92 compared to day 63 for parental cells. JIMT-1 pacR cells exhibited minimal tumorigenicity, forming tumors in only one mouse by day 159. These results suggest that the acquisition of resistance may be coupled with a less proliferative, low-tumorigenic phenotype *in vivo*, potentially reflecting survival-driven adaptations at the expense of aggressive growth. Hormonal receptor expression status in tumor xenografts was evaluated by IHC, revealing a consistent expression pattern (Fig. S3A).

### 3.2. Chromosomal alterations and deleterious variants emerge across multiple pathways in resistant cells

To further investigate the molecular mechanisms underlying the phenotypic changes observed in drug-resistant cells, we performed a comprehensive examination of genomic alterations. This revealed widespread copy number alterations (CNAs) across all cell lines (Fig. 2A). In MDA-MB-231 parental cells, a 2.9 Mb copy number gain at cytoband 17q12 - 17q21.2, encompassing the *TOP2A* gene, was lost in MDA-MB-231 doxR cells. Similarly, large gains in the parental line, including a 69.9 Mb region at 4p16.3 – 4q13.2 involving *SOD3* and a 17.58 Mb region at 4q13.2 – 4q21.3 including deoxycytidine kinase (*DCK*), were reversed to a diploid state in the resistant cells. In JIMT-1 doxR cells, an 11.21 Mb gain at 16p13.12 – 16p12.1 harboring *ABCC1* was observed, while a 101.62 Mb loss spanning 4q13.2 – 4q33 covering *DCK*, *NFKB1*, *METTL14,* and *FGF2* in parental cells was reverted to diploidy. In T-47D doxR cells, a 65.89 Mb loss was detected at 4p15.2 – 4q21.3 involving *SOD3* and *DCK*. Several alterations were shared across resistant cell lines. For example, both MDA-MB-231 pacR and JIMT-1 pacR cells exhibited either loss or reversion to diploidy in a 17.95 Mb region at 6p22.3 – 6p21.31 harboring *ALDH5A1*, *TPMT*, *SOX4,* or *NOTCH4*, and a 3.52 Mb region at 6p21.2 – 6p21.1 containing *APOBEC2*, *CCND3*. Additional information is provided in Supplementary Table S1, offering a valuable foundation for further investigations aimed at unraveling the complete spectrum of genomic changes that underpin resistance to DOX and PAC.

**Fig. 2.**
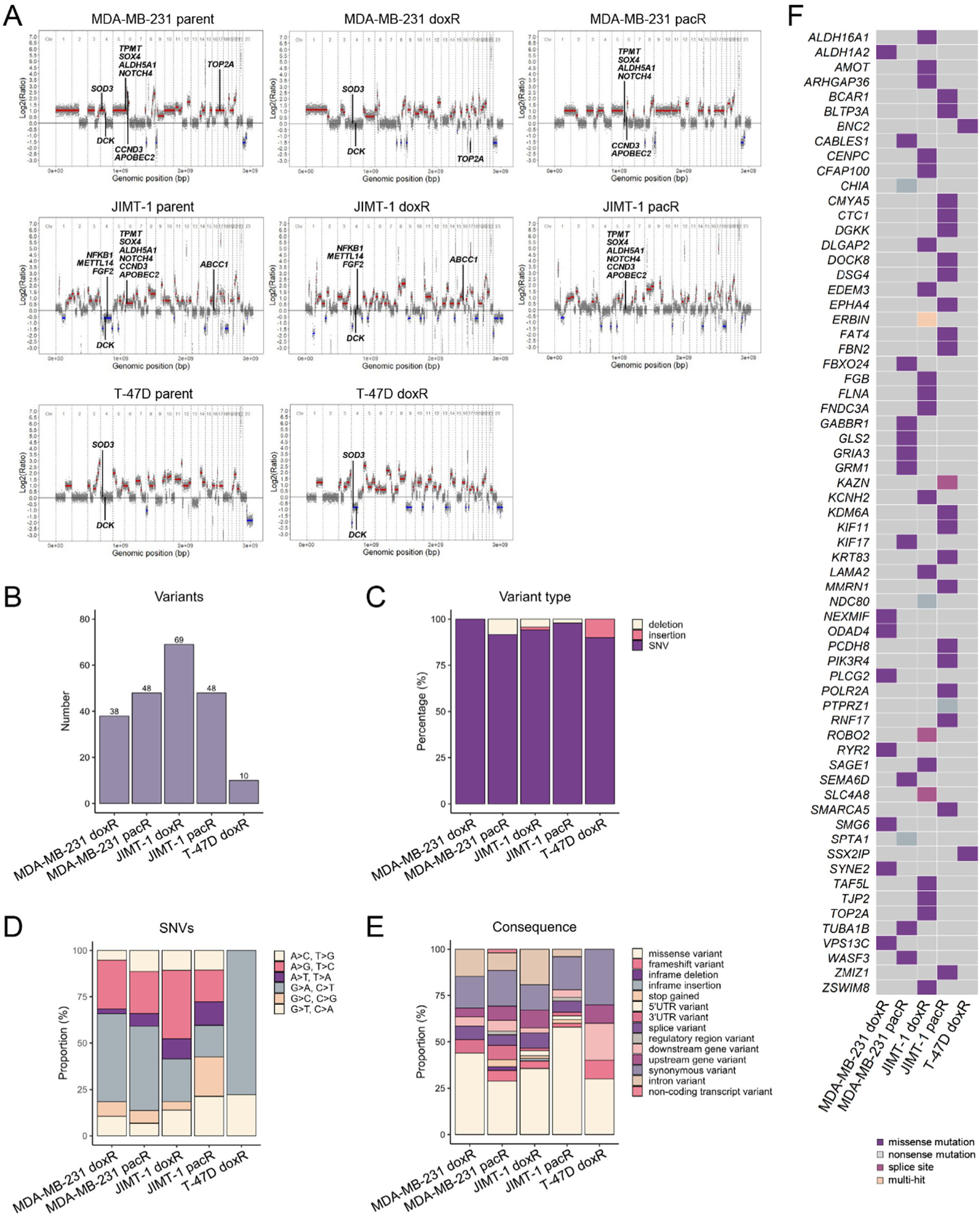
The development of chemoresistance is accompanied by widespread genomic aberrations at both chromosomal and single-nucleotide levels. (A) Chromosomal rearrangements in individual cell lines compared to Female Human Reference DNA. (B) Number of observed mutations in chemotherapy-resistant cells, (C) including a breakdown of individual variant types, (D) single nucleotide variant substitutions, and (E) variant classification. (F) Oncoplot illustrating potentially deleterious variants with a frequency > 0.35 and CADD PHRED score > 20. Parental cell lines were used as a reference. Abbreviations: SNV, single nucleotide variant; UTR, untranslated region.

Whole-exome sequencing was conducted to uncover subtle genomic alterations, such as point mutations and small insertions or deletions, offering a comprehensive view of the genetic landscape linked to chemotherapy resistance. JIMT-1 doxR cells exhibited the highest number of alterations, while T-47D cells showed the fewest. Single nucleotide variants (SNVs), particularly transitions (G>A, C>T), and missense mutations were the most prevalent alterations across resistant cell lines (Fig. 2B-E). Potentially deleterious variants were identified in 67 genes (Fig. 2F). In MDA-MB-231 doxR cells, missense mutations were detected in *ALDH1A2* (c.272G>A, p.G91D), involved in aldehyde metabolism, and *PLCG2* (c.1918C>T, p.P640S), a key regulator of signal transduction. MDA-MB-231 pacR cells harbored mutations in *CABLES1* (c.532G>A, p.E178K), a cell cycle regulator, *TUBA1B* (c.748G>A, p.V250I), which encodes a microtubule isoform, and *GRM1* (c.586A>C, p.T196P), associated with glutamate signaling. JIMT-1 doxR cells exhibited mutations in *TOP2A* (c.2291T>C, p.L764P), the primary target of DOX, and *NDC80* (c.1432A>T, p.K478X), essential for mitotic spindle stability. JIMT-1 pacR cells possessed mutations in *KIF11* (c.593G>A, p.G198D), a kinesin critical for mitotic spindle function, and *SMARCA5* (c.1478C>G, p.P493R), a chromatin remodeling factor. In T-47D doxR cells, mutations were found in *BNC2* (c.2551C>T, p.H851Y), linked to transcriptional regulation, and *SSX2IP* (c.1016C>A, p.T339K), involved in mitotic spindle organization.

### 3.3. Chemotherapy-resistant cells exhibit dysregulated drug resistance pathways

To determine whether the established chemoresistant models recapitulate features observed in clinical drug resistance, we investigated gene expression changes using the RT^2^ Profiler Human Cancer Drug Resistance assay (Fig. 3A, B). The upregulation of *ABCC3* and *RELB* across several cell lines suggests enhanced drug efflux capacity and altered inflammatory signaling. Downregulation of *TOP2A* in MDA-MB-231 doxR and JIMT-1 doxR cells points to potential disruptions in DNA repair, critical to the chemotherapy response. Notably, the loss of a chromosomal region encompassing *TOP2A* in MDA-MB-231 doxR cells may underlie its reduced expression. The consistent downregulation of *PPARG* across multiple resistant cell lines further indicates impaired metabolic and stress response pathways. Of particular interest is the > 170-fold upregulation of *ABCB1* in JIMT-1 pacR cells, indicating a substantial increase in drug efflux likely contributing to PAC resistance. Finally, changes in *FGF2, ABCG2*, and *GSTP1* expression further confirm widespread alterations in drug transport and detoxification processes.

**Fig. 3.**
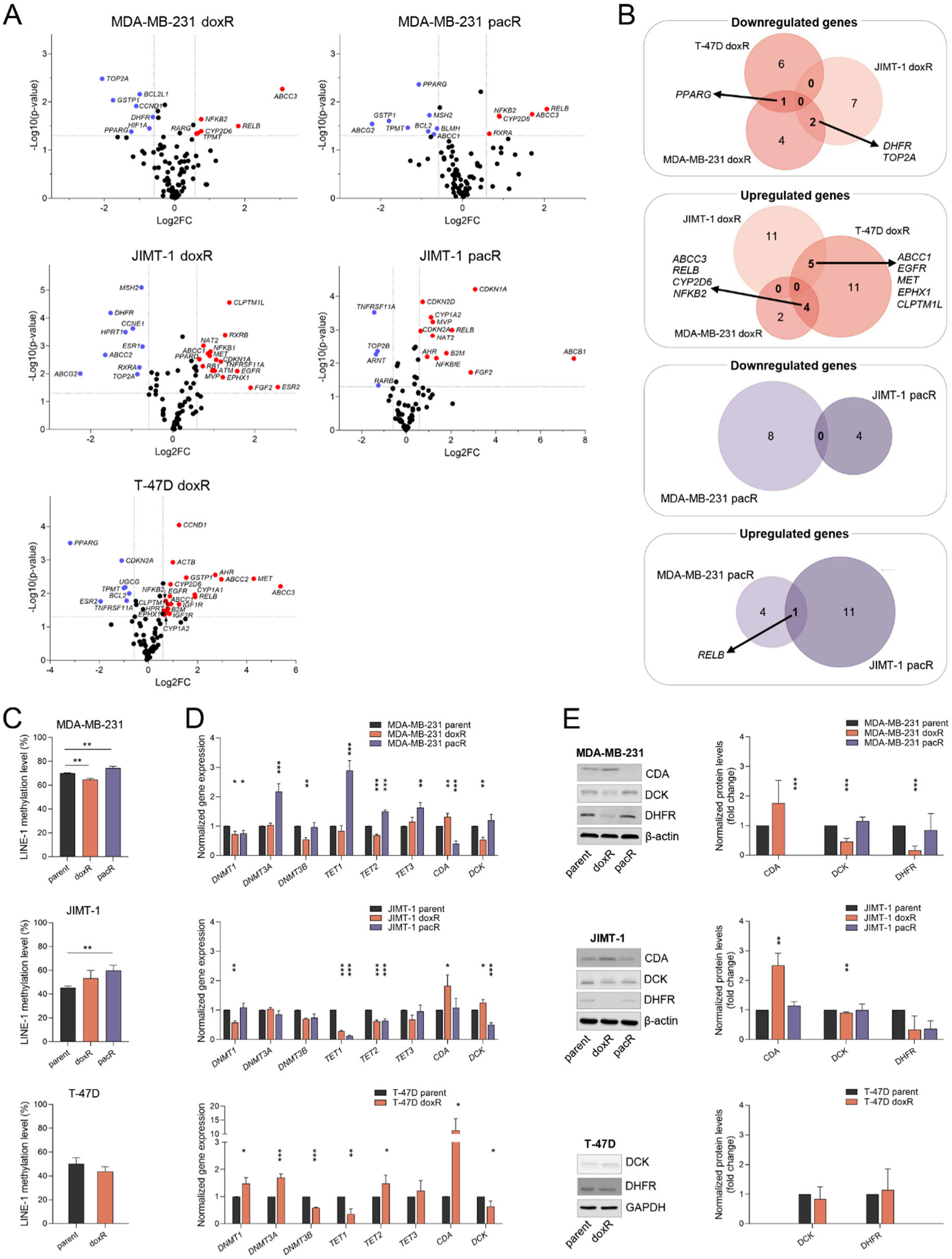
Chemoresistant cells exhibit dysregulated expression of numerous genes involved in drug resistance and DNA methylation pathways, accompanied by changes in global DNA methylation. (A) Volcano plots showing significantly upregulated (red) or downregulated (blue) genes (|FC| > 1.5) in MDA-MB-231, JIMT-1, and T-47D chemotherapy-resistant cells relative to their parental counterparts. Data represent the mean from three independent experiments. (B) Venn diagrams illustrating the overlap of deregulated genes among the individual cell lines. (C) LINE-1 methylation levels in parental and chemotherapy-resistant cells, expressed as mean methylation (%) of three CpG sites within the LINE-1 sequence. (D) Normalized expression of key epigenetic regulators and genes involved in nucleoside metabolism. (E) Quantification of corresponding protein expression by Western blot. All data are presented as mean ± SD from three independent experiments. * p < 0.05, ** p < 0.01, *** p < 0.001.

Given the observed downregulation of dihydrofolate reductase (*DHFR*) in MDA-MB-231 and JIMT-1 doxR cell lines, we sought to investigate its potential impact on nucleotide metabolism and DNA methylation pathways. DHFR is a key enzyme in folate metabolism, and its reduced expression may impair nucleotide biosynthesis by limiting the availability of tetrahydrofolate (THF), a cofactor essential for purine and thymidylate biosynthesis. THF also contributes to one-carbon metabolism, which generates S-adenosylmethionine (SAM), the primary methyl donor for DNA methylation processes. Despite this, LINE-1 analysis revealed divergent global DNA methylation patterns across studied cell lines (Fig. 3C). In MDA-MB-231 doxR cells, decreased global DNA methylation was accompanied by modest downregulation of *DNMT1, DNMT3B,* and *TET2* (Fig. 3D). This trend, however, was not observed in JIMT-1 doxR cells. Conversely, both pacR cell lines displayed significantly increased global DNA methylation, despite inconsistent expression patterns of epigenetic modifiers. The upregulation of all *TET* family genes along with *DNMT3A* in MDA-MB-231 pacR cells suggests potential compensatory or subtype-specific epigenetic remodeling.

Additionally, we observed significant alterations in enzymes involved in nucleotide metabolism, such as cytidine deaminase (CDA) and DCK, which are crucial for DNA repair and replication. Gene encoding CDA, which catalyzes the deamination of nucleosides, was upregulated across all doxR cell lines (Fig. 3D, E), suggesting a shift in cellular metabolism that could influence DNA damage repair mechanisms. Although CDA’s primary role is inactivating nucleoside analogs, its increased expression may reflect a broader change in nucleotide metabolism that could enhance the cells’ ability to handle DNA damage, thereby contributing to DOX resistance. In contrast, DCK, responsible for phosphorylating and activating nucleosides, was downregulated in MDA-MB-231 doxR cells. CDA and DCK expression patterns in pacR cells were more heterogeneous, with no clear correlation between enzyme levels and resistance phenotype.

### 3.4. Development of resistance to PAC in TNBC is accompanied by large-scale DNA methylation reprogramming

To assess the association between DNA methylation and the development of drug resistance in more depth, we compared the methylomic and transcriptomic landscapes of pacR xenografts with those from their parental counterparts. These two models were selected for whole-genome analyses due to their suitability for preclinical evaluation of combination therapies. In contrast, doxR xenografts were excluded owing to their slow tumor growth, which would have limited the feasibility of meaningful comparisons across models.

Our analysis revealed extensive dysregulation of the DNA methylation in pacR xenografts, with 230,052 DMPs, including 188,657 with Δβ > |0.1| and 56,450 with Δβ > |0.3|. Notably, the proportion of hypermethylated CpG islands nearly doubled in pacR xenografts, suggesting a potential impact on gene expression regulation, while methylation patterns in the open sea, shelf, and shore regions remained largely unchanged (Fig. 4A, B, Supplementary Table S2). Consistently, we identified 2,704 DEGs, including 1,647 with a logFC > |1.5|, of which 1,397 were protein-coding genes. Hypermethylated CpG dinucleotides were predominantly enriched in CpG islands, shelves, first exons, exon boundaries, intergenic regions, and transcription start site (TSS) 200, whereas hypomethylation was observed in regions such as the 5′ untranslated regions (5′ UTR) and TSS1500. Consistent with *in vitro* findings, the development of PAC resistance was associated with an overall increase in DNA methylation at CpG sites present on the BeadChip, as reflected by elevated median β values (Fig. 4C) and illustrated in the heatmap of DMPs (Fig. 4D). These observations highlight widespread epigenetic alterations accompanying PAC resistance, which may play a significant role in modulating gene expression.

**Fig. 4.**
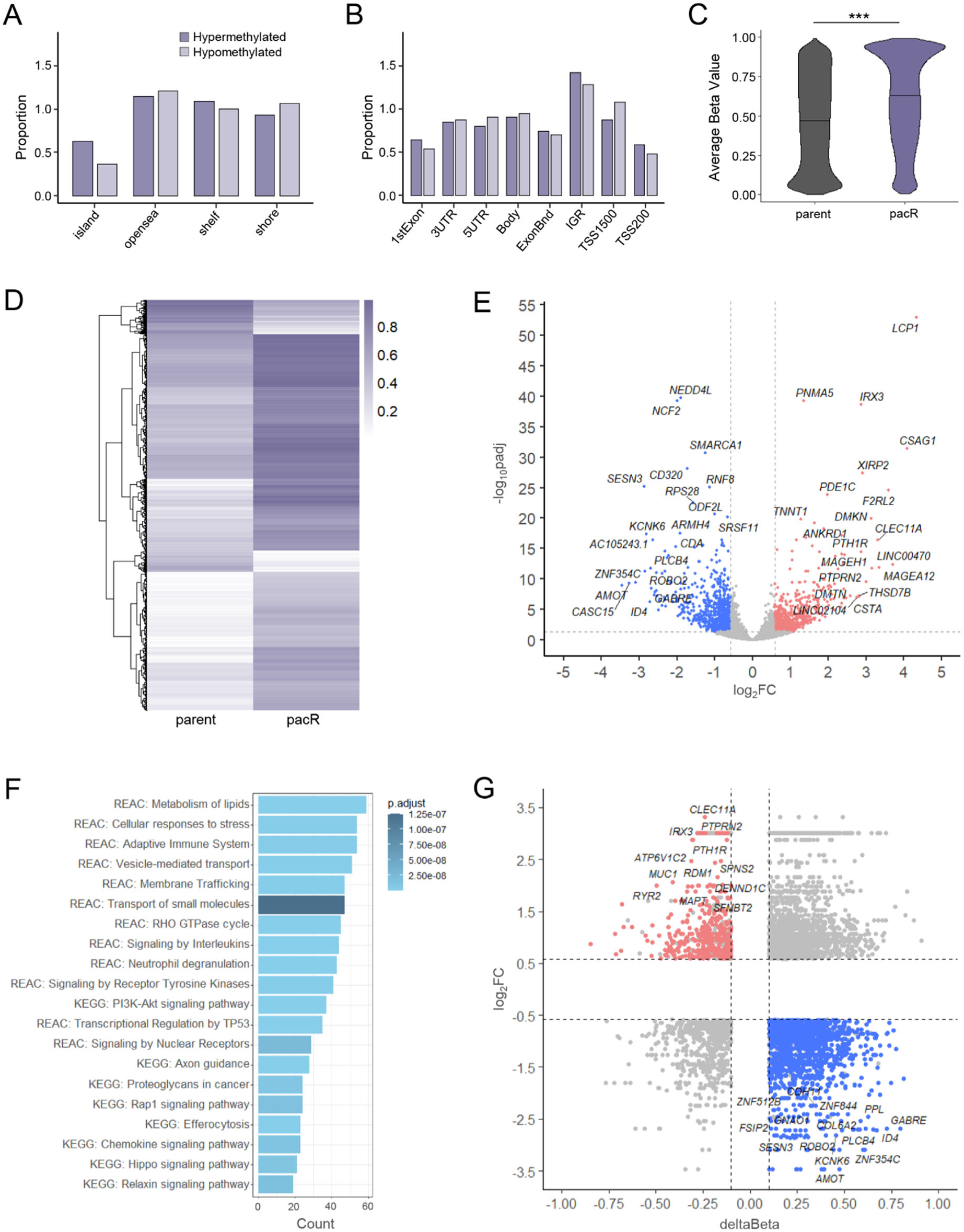
Development of paclitaxel (PAC)-resistance is accompanied by extensive DNA methylation–mediated molecular reprogramming in TNBC xenografts. (A) Distribution of differentially methylated CpGs (DMPs) across CpG islands and (B) genomic features. (C) Violin plots illustrating β values of DMPs (adjusted p-value < 0.05) in parental and pacR xenografts. (D) Heatmap of β values of DMPs (Δβ > |0.1| and adjusted p-value < 0.05). (E) The volcano plot showing gene expression changes in pacR xenografts relative to parental MDA-MB-231 cells, with the top 20 upregulated and downregulated genes highlighted based on log₂ fold change (log₂FC) and statistical significance (adjusted p-value). (F) 20 selected dysregulated signaling pathways associated with paclitaxel resistance. (G) Volcano plot displaying Δβ values of DMPs linked to relevant inversely correlated genes, identified through integrative analysis of gene expression. Abbreviations: FC, fold change; IGR, intergenic region; UTR, untranslated region; TSS, transcription start site; ExonBnd, exon boundaries; IGR, intergenic regions. *** p < 0.01.

Among the five top-upregulated genes in pacR xenografts (Fig. 4E) are *LCP1* and *XIRP2*, *which* regulate actin cytoskeleton dynamics, contributing to immune cell migration and tumor cell motility, *PNMA5*, a tumor-associated antigen, linked to immune evasion, *IRX3*, a homeobox transcription factor, participating in metabolic state and developmental reprogramming, and *CSAG1* involved in the centrosome cycle. The top-downregulated genes include *NEDD4L* and *RNF8*, both E3 ubiquitin ligases with distinct roles in BC. *NEDD4L* primarily functions as a tumor suppressor, regulating pathways such as TGF-β and Wnt by targeting specific substrates for degradation. Conversely, *RNF8* is critical in the DNA damage response, facilitating DNA repair processes. Another downregulated gene, *SMARCA1*, encodes a chromatin remodeler involved in regulating gene expression and maintaining genomic stability. *NCF2*, a component of NADPH oxidase, is essential for reactive oxygen species production and immune defense mechanisms. Additionally, *CD320*, a receptor for vitamin B12 uptake, is critical for cell metabolism and proliferation. Several other findings align with our *in vitro* results, confirming the upregulation of *ABCC3* and the downregulation of *TUBB, CDA, GSTP1, BCL2L,* and *TPMT* (Fig. 3A). Enriched signaling pathways closely align with the top DEGs (Fig. 4F). These included lipid metabolism, cellular response to stress, vesicle-mediated transport, transport of small molecules, transcriptional regulation by TP53, and cancer-related pathways such as PI3K-Akt, Hippo signaling, axon guidance, proteoglycans in cancer, and efferocytosis.

Integrative analysis identified 1,424 hypermethylated and 357 hypomethylated CpG sites located in regulatory regions, including CpG islands, shores, and shelves, affecting 411 hypermethylated and 177 hypomethylated unique genes (Fig. 4G). Pathway analysis confirmed the enrichment of key biological pathways, closely mirroring those observed when all DEGs were considered (Supplementary Table S3). These pathways include cancer signaling, lipid metabolism, apoptosis, and estrogen receptor signaling, all of which play critical roles in tumor progression and therapeutic response. These findings highlight the functional impact of methylation changes on essential regulatory networks and suggest that epigenetic modifications may contribute to both oncogenic processes and potential vulnerabilities in resistant cancer cells.

### 3.5. Decitabine synergizes with chemotherapy in parental TNBC models

Based on our previous findings demonstrating that the DNA demethylating agent DAC sensitizes JIMT-1 cells to DOX (Buocikova et al., 2022a), we extended our investigation to TNBC. We first evaluated the sensitivity of both parental and chemotherapy-resistant MDA-MB-231 cells to DAC. While DAC treatment did not induce significant differences in cell viability between parental and doxR cells, pacR cells showed markedly reduced sensitivity, with a 61-fold increase in the IC_50_ value (Fig. 5A).

**Fig. 5.**
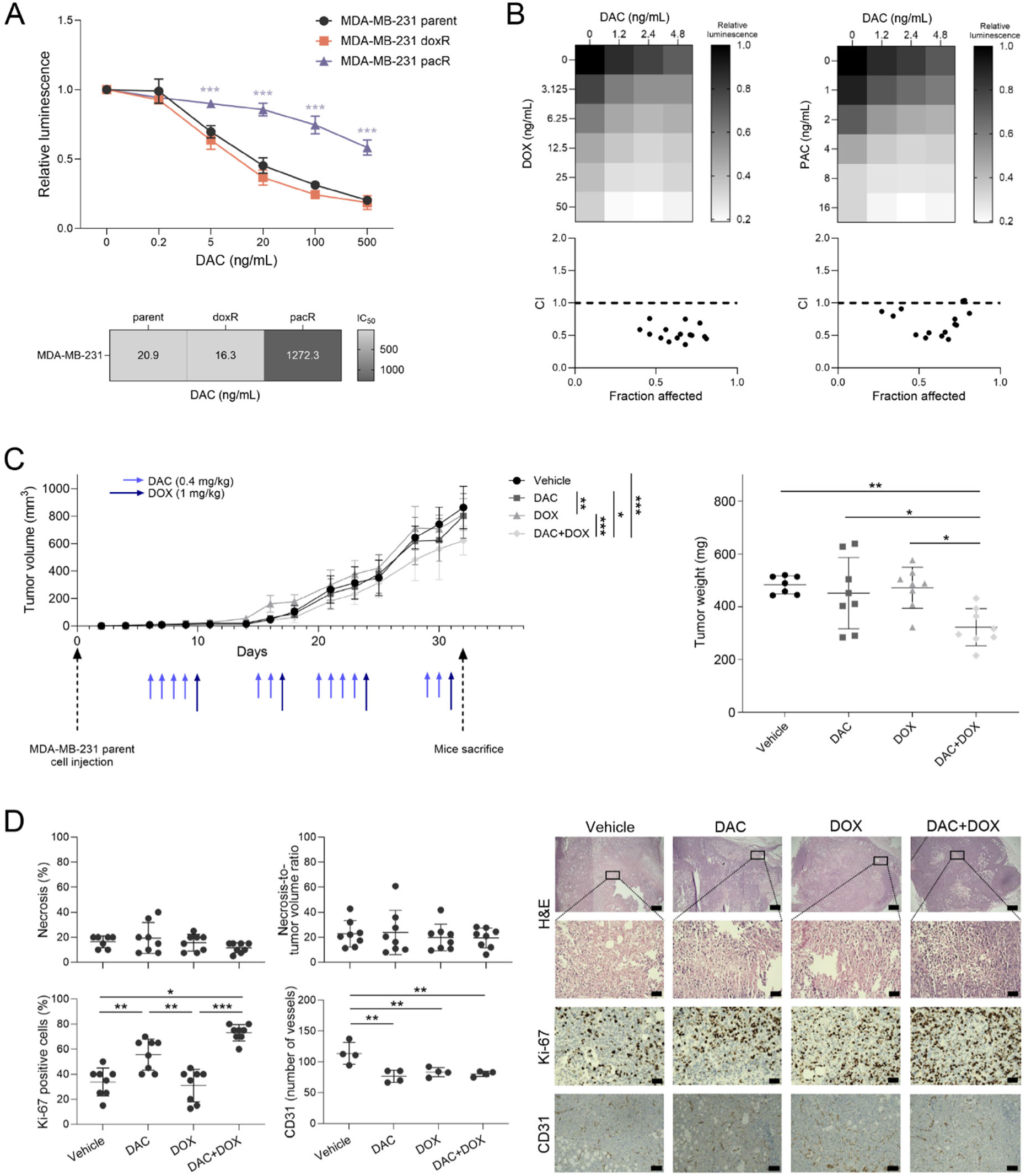
Doxorubicin (DOX) synergizes with decitabine (DAC) in parental MDA-MB-231 cells, leading to reduced tumor growth, alongside increased Ki-67 expression and decreased vascularization. (A) Cell viability of MDA-MB-231 parental, doxorubicin-resistant (doxR), and paclitaxel-resistant (pacR) cells following a 6-day treatment with DAC relative to the untreated controls (set to 1). Statistical significance was determined relative to DAC-treated parental cells. IC_50_ values for DAC in MDA-MB-231 parental, doxR, and pacR cells determined by CalcuSyn software. (B) Viability of MDA-MB-231 parental cells after monotherapy or combination treatment with DAC and DOX or PAC, relative to untreated controls (set to 1). Data are expressed as mean ± SD from three independent experiments. Combination index (CI) values for DAC and DOX or DAC and PAC treatments were calculated using CalcuSyn software. CI < 1 indicates synergy effect, CI = 1 additive effect, and CI > 1 antagonism. (C) MDA-MB-231 parental tumor volumes over time with an indication of the therapeutic scheme and tumor weights for each treatment group. (D) Quantitative evaluation of treatment-induced changes in tumor necrosis (hematoxylin and eosin, H&E), Ki-67, and vessel quantification based on CD31 immunostaining. Horizontal lines indicate the mean. For CD31, each dot represents the average number of blood vessels counted across seven distinct regions of interest. The representative immunohistochemical images of each marker captured at 25× (H&E, 400µm scale bar), 200× (H&E, Ki-67, 50 µm scale bar) and 100× magnification (CD31, 100 µm scale bar) are shown on the right. *p < 0.05, **p < 0.01, ***p < 0.001.

Next, we assessed the efficacy of DAC-based combination therapy in parental MDA-MB-231 cells. Combining DAC with DOX or PAC significantly reduced cell viability compared to monotherapy at most tested concentrations (Fig. 5B). The DAC and DOX combination showed a consistent synergistic effect (CI < 1), while the DAC and PAC combination also exhibited synergy, except for two concentrations where the effect was additive (CI = 1).

To evaluate these findings *in vivo*, mice bearing xenografts were treated with low doses of DAC (0.4 mg/kg), DOX (1 mg/kg), or their combination, following the treatment schedules outlined in Fig. 5C. The combination therapy resulted in significant reduction in both tumor volumes and weight (Fig. 5C). Intriguingly, DAC alone and more prominently in combination with DOX, induced Ki-67 expression, while all treatment groups showed impaired tumor vascularization, as evidenced by reduced CD31 staining in tumor tissues (Fig. 5D).

### 3.6. Combination therapy is more efficient when using agents unrelated to the resistance-inducing drug

To evaluate the efficacy of combination therapy in chemotherapy-resistant models, we initially assessed the potential of DAC to augment the cytotoxic effects of chemotherapeutic agents *in vitro*. In MDA-MB-231 doxR cells, DAC combined with DOX significantly reduced cell viability compared to DOX alone (Fig. 6A), primarily resulting in an additive effect. In contrast, the combination of DAC and PAC predominantly exhibited a synergistic effect. For MDA-MB-231 pacR cells, DAC concentrations were adjusted to 10 - 30 ng/mL (corresponding to IC_20_) due to their reduced sensitivity. The combination of DAC and DOX significantly decreased cell viability compared to DOX alone, with synergy observed at most concentrations (Fig. 6B). The DAC and PAC combination demonstrated higher CI values than the DAC and DOX, yielding antagonistic effects in multiple cases. These results suggest that DAC is more effective when combined with chemotherapy agents to which cells have not developed resistance.

**Fig. 6.**
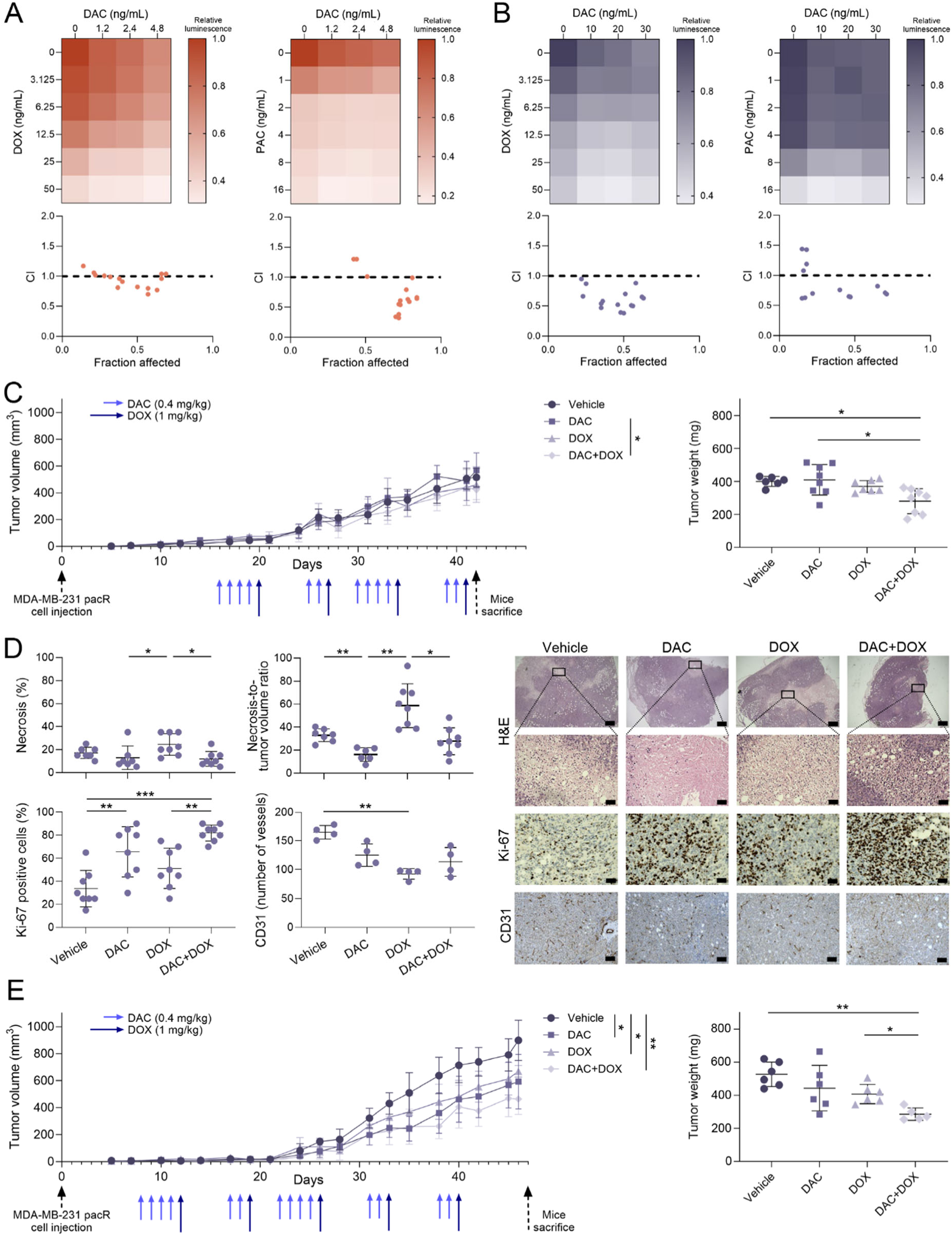
Decitabine enhances the efficacy of chemotherapy agents unrelated to the resistance-inducing drug and improves outcomes in MDA-MB-231 paclitaxel-resistant (pacR) xenografts under an extended treatment regimen. (A) Cell viability of MDA-MB-231 doxorubicin-resistant (doxR) and (B) pacR cells following monotherapy or combined treatment with DAC and chemotherapy (DOX or PAC), normalized to untreated controls (set to 1). Data represent mean ± SD from three independent experiments. Combination index (CI) values were calculated using CalcuSyn software. CI < 1 indicates synergy, CI = 1 additive effect, and CI > 1 antagonism. (C) Treatment-induced changes in pacR tumor volumes and tumor weights at study endpoint, shown alongside the treatment schedule. (D) Quantitative evaluation of differences in tumor necrosis (hematoxylin and eosin, H&E), Ki-67, and vessel quantification based on CD31 immunostaining. Horizontal lines indicate the mean. For CD31, each dot represents the average number of blood vessels counted across seven distinct regions of interest. Representative immunohistochemical images of each marker, captured at 25× (H&E, 400µm scale bar), 200× (H&E, Ki-67, 50 µm scale bar) and 100× magnification (CD31, 100 µm scale bar), are shown on the right. (E) Assessment of therapeutic efficacy following prolonged mono- and combination therapy shown alongside the treatment schedule. *p < 0.05, **p < 0.01, ***p < 0.001.

Following the result of *in vitro* testing, we proceeded to test the combination therapy in pacR xenografts. The MDA-MB-231 doxR model was excluded as explained earlier due to slow tumor growth (Fig. S3B). To ensure valid molecular comparisons, the same treatment regimen was applied across both parental and pacR models, with a ten-day delay in the pacR group to account for slower tumor progression (Fig. 6C). Despite this adjustment, untreated pacR tumors remained nearly half the size of parental tumors by the study endpoint. While combination treatment did not lead to significant changes in tumor volume, it did result in a moderate reduction in tumor weight. Similar to observations in parental tumors, DAC treatment notably increased Ki-67 expression in the pacR cohort. Moreover, Ki-67 levels inversely correlated with tumor necrosis (r = –0.55, p = 0.002 for absolute necrosis; r = – 0.46, p = 0.013 for necrosis normalized to tumor volume) (Fig. 6D). Only DOX monotherapy showed a significant anti-angiogenic effect in the pacR group.

Recognizing that the relatively short treatment window may have limited measurable responses in slow-growing resistant tumors, we extended the duration in a subsequent experiment (Fig. 6E). Under this adjusted regimen, combination therapy significantly decreased both tumor volume and weight, supporting its therapeutic potential in the pacR model.

### 3.7. Decitabine normalizes aberrant methylation changes in paclitaxel-resistant tumor xenografts

The primary goal of this study was to evaluate DAC’s capacity to normalize aberrant DNA methylation patterns in drug-resistant tumors. DAC treatment significantly altered the methylation landscape in both parental and pacR tumor xenografts. In parental xenografts, we identified 100,449 DMPs, predominantly hypomethylated (98,782), with Δ β values ranging from 0.020 to 0.528 (median 0.096). Hypermethylated CpGs showed Δ β values ranging from 0.023 to 0.260, with a median of 0.070. In contrast, pacR xenografts exhibited over twice as many DMPs (213,066), including 207,448 hypomethylated and 5,618 hypermethylated CpGs. The Δ β values for hypomethylated CpGs ranged from 0.019 to 0.574, with a median of 0.124, and for hypermethylated CpGs, the range was 0.019 to 0.365, with a median of 0.079. Hypomethylation was mainly observed in gene bodies, 3’ UTRs, and exon boundary regions, while hypermethylation was more frequent in the first exon and intergenic regions (Fig. 7A, B, Supplementary Table S2). Interestingly, pacR xenografts showed a higher proportion of hypermethylated CpGs in CpG islands, shores, and TSS-proximal regions. Overall, DAC induced a global reduction in β values, reflecting widespread hypomethylation (Fig. 7C).

**Fig. 7.**
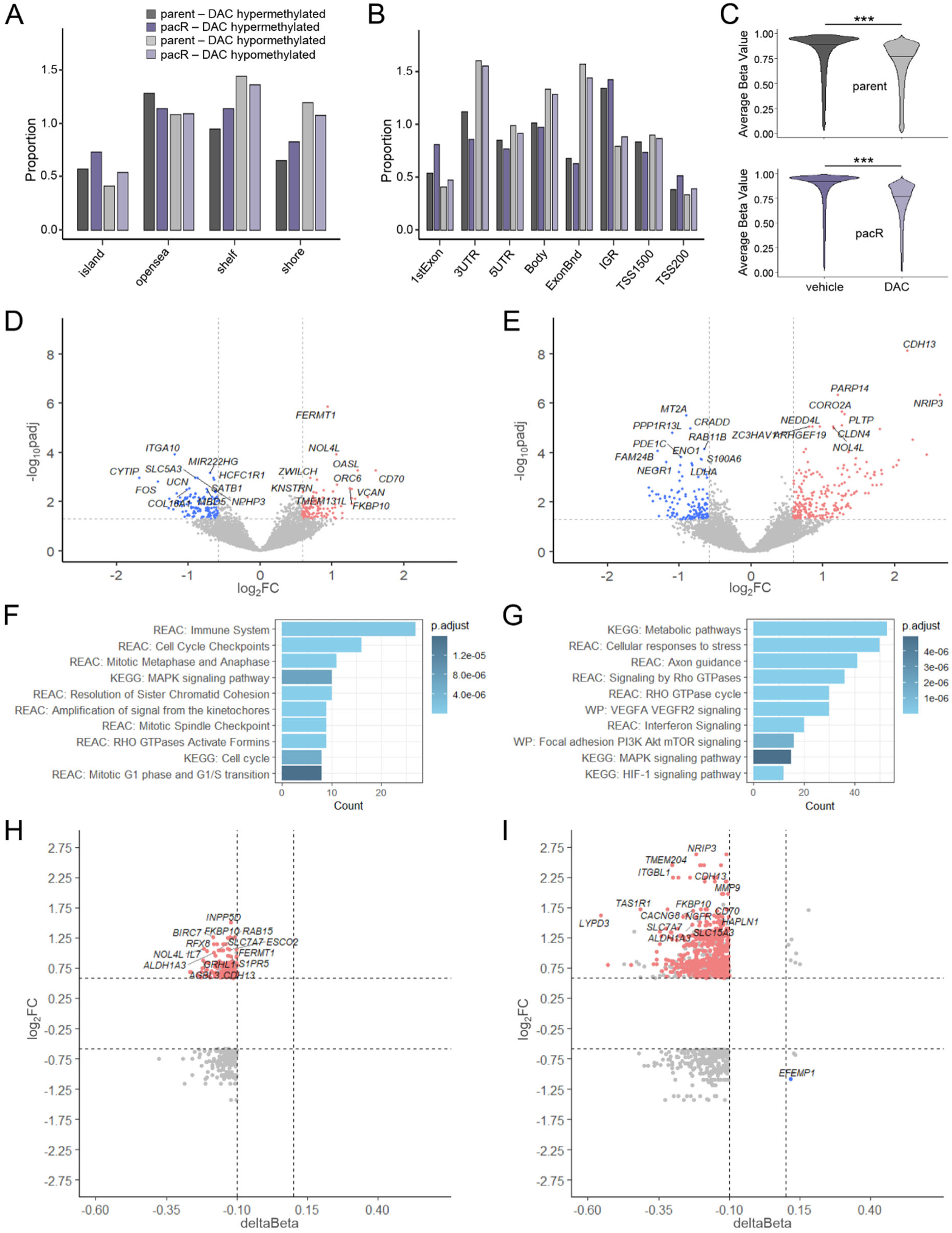
Decitabine (DAC) reduces DNA methylation in both parental and paclitaxel-resistant (pacR) MDA-MB-231 tumor xenografts. (A) Distribution of differentially methylated CpGs across CpG islands and (B) genomic features. (C) Violin plots illustrating DNA methylation changes following DAC treatment. (D) Volcano plot depicting gene expression changes in parental and (E) pacR xenografts after DAC treatment, highlighting the top 20 upregulated and downregulated genes based on log₂ fold change (log₂FC) and statistical significance (adjusted p-value). (F) 10 selected enriched signaling pathways following DAC treatment in parental and (G) pacR xenografts. (H) Volcano plot showing Δ β values of differentially methylated CpGs associated with relevant genes, identified through integrative analysis of gene expression (log₂FC) in parental and (I) pacR xenografts. Abbreviations: FC, fold change; IGR, intergenic region; UTR, untranslated region; TSS, transcription start site; ExonBnd, exon boundaries; IGR, intergenic regions. *** p < 0.01.

Gene expression changes following DAC were more modest than those observed during the development of drug resistance. In parental tumors, DAC upregulated 116 and downregulated 89 protein-coding genes, primarily involved in cell adhesion, migration, and mitotic regulation. Key upregulated genes included *FERMT1, ZWILCH*, and *KNSTRN,* which regulate integrin signaling and mitotic fidelity, while downregulated genes such as *ITGA10* and *CYTIP*, *FOS*, and chromatin remodelers *SATB1* and *MBD5* suggest impaired adhesion, migration, and proliferation (Fig. 7D). In pacR tumors, DAC affected a larger set of genes, 223 upregulated and 120 downregulated, including *CDH13, CLDN4*, and *CORO2A,* implicating cytoskeletal remodeling and tight junction integrity. Downregulation of metabolic and apoptotic regulators, including *ENO1*, *LDHA, CRADD*, and *S100A6,* indicates potential shifts in cell survival and metabolic adaptation (Fig. 7E). Pathway analysis in parental xenografts revealed enrichment in cell cycle regulation, whereas pacR tumors showed activation of cancer-associated pathways, metabolic reprogramming, Rho GTPase, and HIF-1 signaling (Fig. 7F, G).

Integrative analysis identified DAC-induced upregulation of 68 genes (139 CpGs) in parental and 161 genes (592 CpGs) in pacR tumors, predominantly in regulatory regions (Fig. 7H, I). Of these, 22 genes overlapped with loci that became hypermethylated during resistance acquisition. DAC reactivated tumor suppressors such as *FHL1, NEDD4L*, *TSPYL5*, and *CDH13*, but also pro-oncogenic genes including *SRC, MMP16*, *PARP14,* and cancer stem cell marker *ALDH1A3*. These findings suggest that while DAC partially reverses resistance-associated DNA hypermethylation and reactivates silenced tumor suppressors, it may concurrently promote the expression of oncogenic pathways. This dual effect underscores the complexity of epigenetic therapy and highlights the need for selective strategies to maximize benefits.

## 4. Discussion

Systemic treatment, encompassing endocrine therapy, targeted therapy, and chemotherapy, remains a cornerstone of BC management. While patients with hormone receptor-positive or HER2-positive subtypes often benefit from endocrine or targeted therapies, chemotherapy is the primary option for those with advanced-stage disease and TNBC. However, the emergence of therapeutic resistance poses a significant challenge to treatment efficacy, contributing to disease progression and poor clinical outcomes. Despite advances in understanding certain resistance mechanisms, the multifactorial nature of this phenomenon necessitates further investigation.

In this study, we developed robust *in vitro* and *in vivo* models to investigate chemotherapy resistance to two widely used chemotherapeutics, the topoisomerase II inhibitor DOX and the microtubule inhibitor PAC. The derived resistant cell lines exhibited 4.7-to 7.8-fold increase in IC_50_ values, reflecting clinically relevant resistance levels. Although some models report extreme resistance (over 100-fold), evidence from patient-derived cells suggests that more moderate shifts, such as those observed here, more accurately reflect resistance seen in clinical settings (McDermott et al., 2014). Phenotypic characterization revealed slower proliferation and increased doubling times in resistant cells, consistent with clinical observations. Transcriptomic analyses of tumor biopsies before and after neoadjuvant chemotherapy in BC patients have shown that high expression of proliferative and immune-related genes correlates with better therapeutic response, while post-treatment tumors frequently show downregulation of these genes, possibly as an adaptive mechanism (Hoogstraat et al., 2022). Alterations in cell cycle regulation within chemotherapy-resistant cancer cells may further contribute to the observed reduction in cancer cell proliferation.

Acquisition of chemotherapy resistance is accompanied by the emergence of genomic alterations (Kazan et al., 2025; Shadeo and Lam, 2006). Our genomic analysis revealed extensive CNAs in resistant cell lines, indicating ongoing genomic instability under therapeutic pressure. Notably, loss of the *TOP2A*-containing region in MDA-MB-231 doxR cells may impair DOX efficiency, given that *TOP2A* is a direct target of the drug. Conversely, the copy number gain at the locus containing the drug efflux transporter gene *ABCC1* in JIMT-1 doxR cells may contribute to multidrug resistance (Tsyganov et al., 2022). Other recurrent CNAs included alterations in *NOTCH4* and *CCND3*, genes implicated in tumor survival and progression (Tian et al., 2023; Wang et al., 2024), suggesting convergent evolution across different resistance pathways. In addition to CNAs, we identified potentially pathogenic point mutations. A missense mutation in *TOP2A* c.2291T>C (p.L764P) was found in JIMT-1 doxR cells. Given DOX’s mechanism of action, mutations in *TOP2A* can directly mediate resistance (Grimsley et al., 2024; Jado et al., 2024). Decreased *TOP2A* expression has also been linked to resistance in numerous models (Burgess et al., 2008; Kumar et al., 2023; Tegze et al., 2012). In MDA-MB-231 pacR cells, a *TUBA1B* c.748G>A (p.V250I) was identified, potentially altering microtubule dynamics. Elevated *TUBA1B* expression has similarly been implicated in PAC resistance in other cancers (Lu et al., 2013). Upregulation of *ABCB1* in JIMT-1 pacR cells supports previous findings that associate overexpression of ABC family members with resistance to taxanes and other chemotherapeutics (Nemcova-Furstova et al., 2016; Wang et al., 2014). Additionally, the observed upregulation of *RELB* in MDA-MB-231, JIMT-1, and T47-D resistant cells aligns with its established role in promoting resistance to both chemotherapy and endocrine therapy. Notably, silencing *RELB* has been shown to sensitize cancer cells to treatment by inducing cell cycle arrest (Wang et al., 2020a; Xu et al., 2023; Zhou et al., 2018). In addition, we observed *CDA* upregulation across all derived doxR cells, while *PPARG* was downregulated in multiple resistant cell lines, highlighting these genes as potential candidates for future investigation. Moreover, previous studies have shown that chemotherapy-induced *CDA* upregulation renders cells susceptible to additional therapies such as capecitabine or its metabolite 5’-deoxy-5-fluorocytidine, suggesting a potential therapeutic strategy for *CDA*-overexpressing cancer cells (Gao et al., 2021; Karatkevich et al., 2024).

Downregulation of *DHFR* in our doxR cells suggests a shift in the cellular response to chemotherapeutic stress. Deregulation of methionine metabolism, driven by metabolic reprogramming and alterations in one-carbon metabolism and SAM bioavailability, has previously been linked to DNA hypomethylation in transposable element regions in taxane-resistant BC cells (Deblois et al., 2018). In contrast, our pacR cells exhibited increased LINE-1 methylation, as well as higher β values in MDA-MB-231 pacR tumor xenografts, similar to Gomez-Miragaya et al., who reported the accumulation of DNA methylation in docetaxel-resistant TNBC PDXs, revealing a TNBC-specific resistance signature (Gomez-Miragaya et al., 2019). Moreover, our findings align with previous study showing that PAC treatment induces LINE-1 methylation through DNMT3A binding to the inner gene region of LINE-1, leading to its increased expression (Wang et al., 2020b). In this context, combining hypo- and hypermethylating agents, DAC and SAM, showed reduced tumor burden and metastasis in MDA-MB-231 xenografts, supporting the potential of dual epigenetic targeting strategies (Mahmood et al., 2020).

The role of epigenetic remodeling in chemotherapy resistance is further supported by findings in patient-derived models. Protein levels of DNMTs have been shown to correlate with DAC response in organoids generated from patient-derived xenografts of both chemotherapy-sensitive and -resistant TNBCs, highlighting DNMT expression as a potential predictive biomarker of therapeutic efficacy (Yu et al., 2018). However, DNMT expression in our study showed variable patterns, providing limited support for these associations.

Consistent with our previous findings (Buocikova et al., 2022a), we observed a synergistic effect of combining DAC with either DOX or PAC in MDA-MB-231 parental cells. This aligns with prior studies showing that DAC and DOX-loaded nanoparticles enhanced the elimination of cancer stem-like cells and increased apoptosis in MDA-MB-231 xenografts (Li et al., 2015). DAC has also been reported to sensitize TNBC cells to cisplatin through *NOXA* upregulation (Nakajima et al., 2022) and to promote pyroptosis via gasdermin E reactivation in PAC-resistant MCF7 cells (Gong et al., 2023). Moreover, DAC combined sequentially with DOX has been shown to induce G2/M arrest in resistant MCF7 cells (Vijayaraghavalu et al., 2013), and similar sensitization effects have been noted in other cancer models (Shin et al., 2013). However, in our current study, a key consequence of DAC treatment was increased Ki-67 expression in both parental and pacR tumors. This proliferation shift may temporarily increase their susceptibility to cell cycle-dependent therapies like DOX.

Transcriptomic analyses of DAC-treated xenografts revealed distinct biological responses between parental and resistant tumors. Consistent with previous studies, parental xenografts exhibited upregulation of pathways involved in cell cycle regulation, DNA damage response, and immune activation. DAC has also been previously shown to disrupt 3D chromatin architecture and restore ER-regulated transcription in BC PDXs through global demethylation (Achinger-Kawecka et al., 2024), underscoring its potential in epigenetically targeted therapies. However, in pacR xenografts, DAC treatment was associated with the enrichment of pro-survival and metabolic pathways, including PI3K-Akt-mTOR and HIF-1 signaling, which may contribute to adaptive resistance and limit long-term therapeutic efficacy.

While our models offer valuable insights into resistance mechanisms, some limitations must be acknowledged. Established cell lines may harbor pre-existing genomic aberrations that are not representative of primary tumors, potentially affecting generalizability (Tsuji et al., 2010). *In vivo* experiments were also constrained by the toxicity profile of DOX in SCID/bg mice, limiting our ability to assess clinically relevant doses. Finally, although we identified numerous candidate alterations, further functional validation is needed to confirm their causal role in resistance and therapeutic response.

## 5. Conclusion

Our study provides a comprehensive analysis of the molecular mechanisms underlying chemotherapy resistance in BC, revealing key genomic, transcriptomic, and epigenetic alterations associated with drug-resistant phenotypes. By developing robust *in vitro* and *in vivo* models, we demonstrated clinically relevant resistance patterns to DOX and PAC, characterized by reduced proliferation, altered cell cycle regulation, and extensive genomic instability. CNAs, mutations in drug target genes, and dysregulated expression of key resistance-associated genes highlight diverse yet converging adaptive mechanisms in resistant cells. Notably, DNA methylation profiling uncovered epigenetic reprogramming in resistant tumors, with DAC treatment inducing distinct pathway modulations in parental and pacR xenografts. While DAC enhanced chemotherapy sensitivity in some models, its ability to activate survival pathways in pacR tumors suggests the need for strategic treatment sequencing. Together, our findings provide new insights into the complexity of chemotherapy resistance and offer a foundation for refining therapeutic approaches in treatment-refractory BC.

## CRediT authorship contribution statement

**Lenka Trnkova** – Writing - original draft, Writing - Review & Editing, Investigation, Conceptualization, Funding Acquisition, Formal Analysis, Visualization. **Monika Burikova** – Investigation, Writing - Review & Editing. **Andrea Soltysova** – Investigation, Formal Analysis, Writing - Review & Editing. **Andrej Ficek** – Investigation, Formal Analysis. **Jana Plava** – Investigation. **Andrea Cumova** – Methodology. **Lucia Rojikova** – Methodology. **Kristina Jakic** – Methodology. **Eva Sedlackova** – Methodology. **Boris Tichy** – Resources, Investigation. **Vojtech Bystry** – Resources, Formal Analysis. **Florence Busato** – Investigation, Formal Analysis. **Yimin Shen** – Formal Analysis. **Miroslava Matuskova** – Investigation, Writing - Review & Editing. **Lucia Kucerova** – Funding Acquisition, Writing – Review & Editing. **Jorg Tost** – Resources, Investigation, Formal Analysis, Writing – Review & Editing. **Svetlana Miklikova** – Supervision, Writing – Review & Editing, Project Administration. **Marina Cihova** – Supervision, Project Administration, Funding Acquisition, Writing - Review & Editing. **Verona Buocikova** – Writing - review & editing, Visualization. **Bozena Smolkova** – Supervision, Conceptualization, Writing – Original Draft, Review & Editing.

## Declaration of generative AI and AI-assisted technologies in the writing process

During the preparation of this work the author(s) used ChatGPT-4o mini in order to improve the readability and language of the manuscript. After using this tool/service, the author(s) reviewed and edited the content as needed and take(s) full responsibility for the content of the published article.

## Acknowledgements

We would like to thank the Slovak Cancer Research Foundation for their continuous help and support. Graphical abstract and Supplementary Figure S2A were created with BioRender.com.

## Data Availability

Transcriptomic data are available at Gene Expression Omnibus under accession GSE293933. Whole exome sequencing data were submitted to Sequence Read Archive under accession PRJNA1250986. Additional data generated or analyzed during this study are included in this article and its Supplementary Material files.

