## Supplementary Material for "Development of Paclitaxel Resistance in Triple-Negative Breast Cancer Is Associated with Extensive DNA Methylation Changes That Are Partially Reversed by Decitabine"

**Fig. S1:** Schematic overview of mono- and combined therapy with decitabine (DAC) and doxorubicin (DOX) in MDA-MB-231 parental and chemotherapy-resistant xenograft mouse models.

**Fig. S2:** Derivation of chemotherapy-resistant breast cancer cell lines and their molecular characterization.

**Fig. S3:** Expression of hormone receptors and HER2 in cell line-derived xenografts, and evaluation of monotherapy and combination therapy in a pilot *in vivo* doxorubicin-resistant (doxR) model.

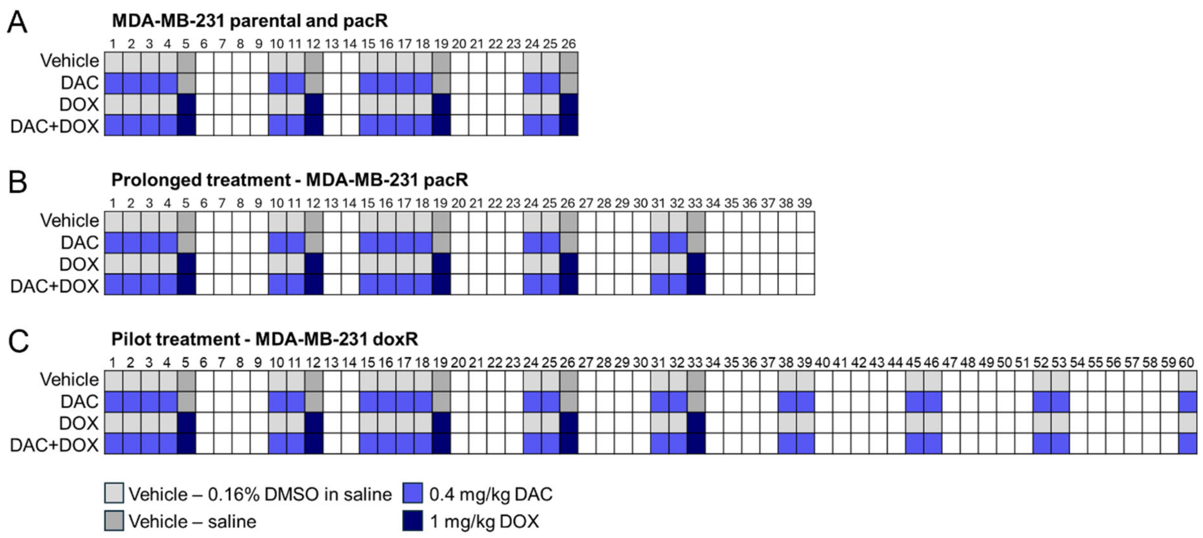


**Fig. S1.** Schematic overview of mono- and combined therapy with decitabine (DAC) and doxorubicin (DOX) in MDA-MB-231 parental and chemotherapy-resistant xenograft mouse models. (A) 26-day treatment regimen in mice bearing MDA-MB-231 parental and paclitaxel-resistant (pacR) xenografts. (B) Extended 39-day treatment in mice with MDA-MB-231 pacR xenografts. (C) Pilot 60-day treatment in mice implanted with MDA-MB-231 doxorubicin-resistant (doxR) xenografts. Abbreviations: DMSO, dimethyl sulfoxide

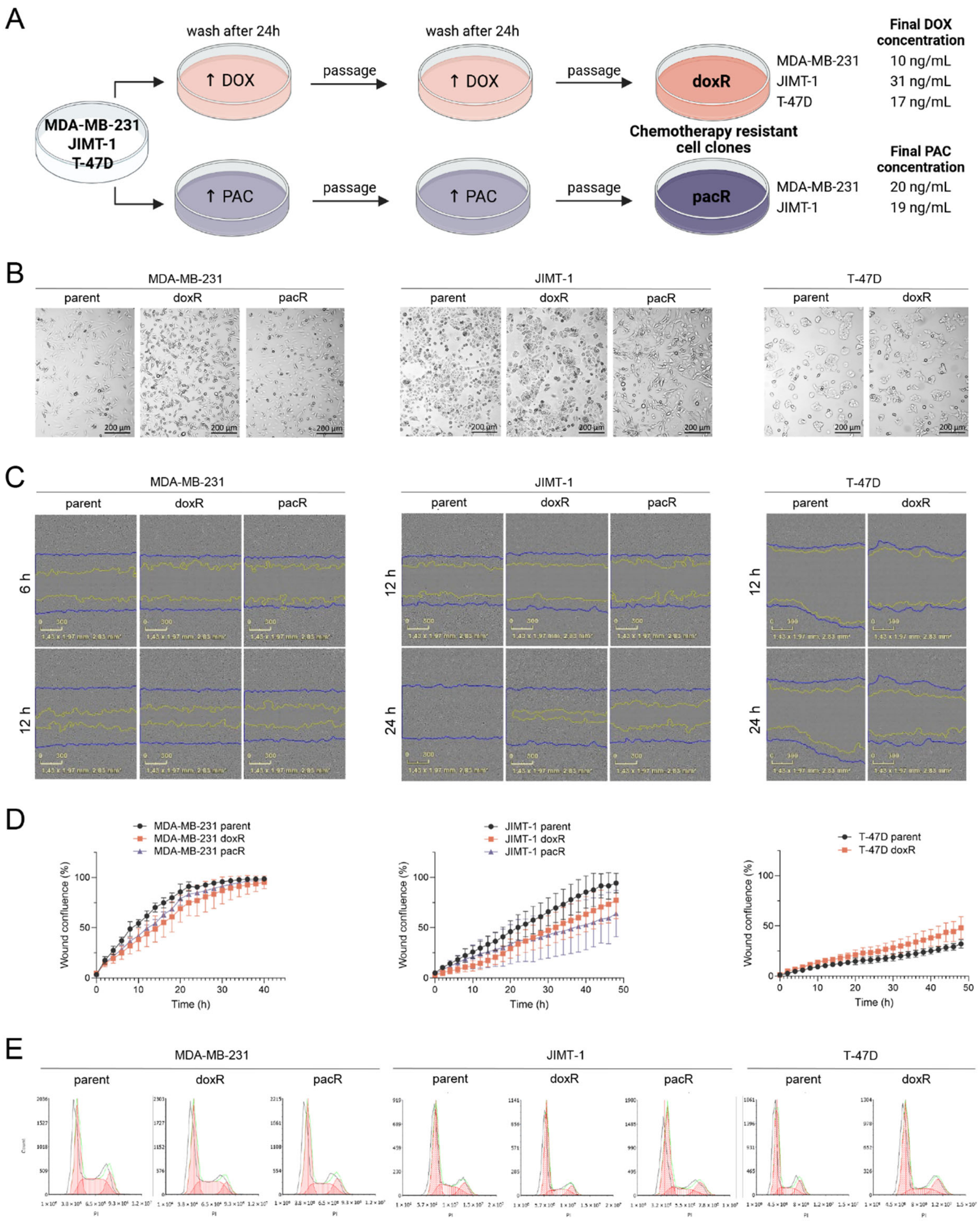


**Fig. S2.** Derivation of chemotherapy-resistant breast cancer cell lines and their molecular characterization. (A) Schematic representation of the experimental strategy for generating doxorubicin-resistant (doxR) and paclitaxel-resistant (pacR) cell lines, including the highest drug concentration applied. (B) Phase-contrast microscopy images of parental, doxR, and pacR cells at 100× magnification (200 µm scale bar). (C) Representative images of wound healing assay at 6 h and 12 h (MDA-MB-231) or 24 h and 48 h (JIMT-1 and T-47D). (D) Migration of parental and chemotherapy-resistant breast cancer cells shown as a percentage of wound confluence over time. Analysis was performed using the IncuCyte$^{TM}$ ZOOM system. Data represent mean ± SD from three independent experiments; no statistically significant changes were observed. (E) Representative images of cell cycle analysis of parental and chemotherapy-resistant cells by flow cytometry.

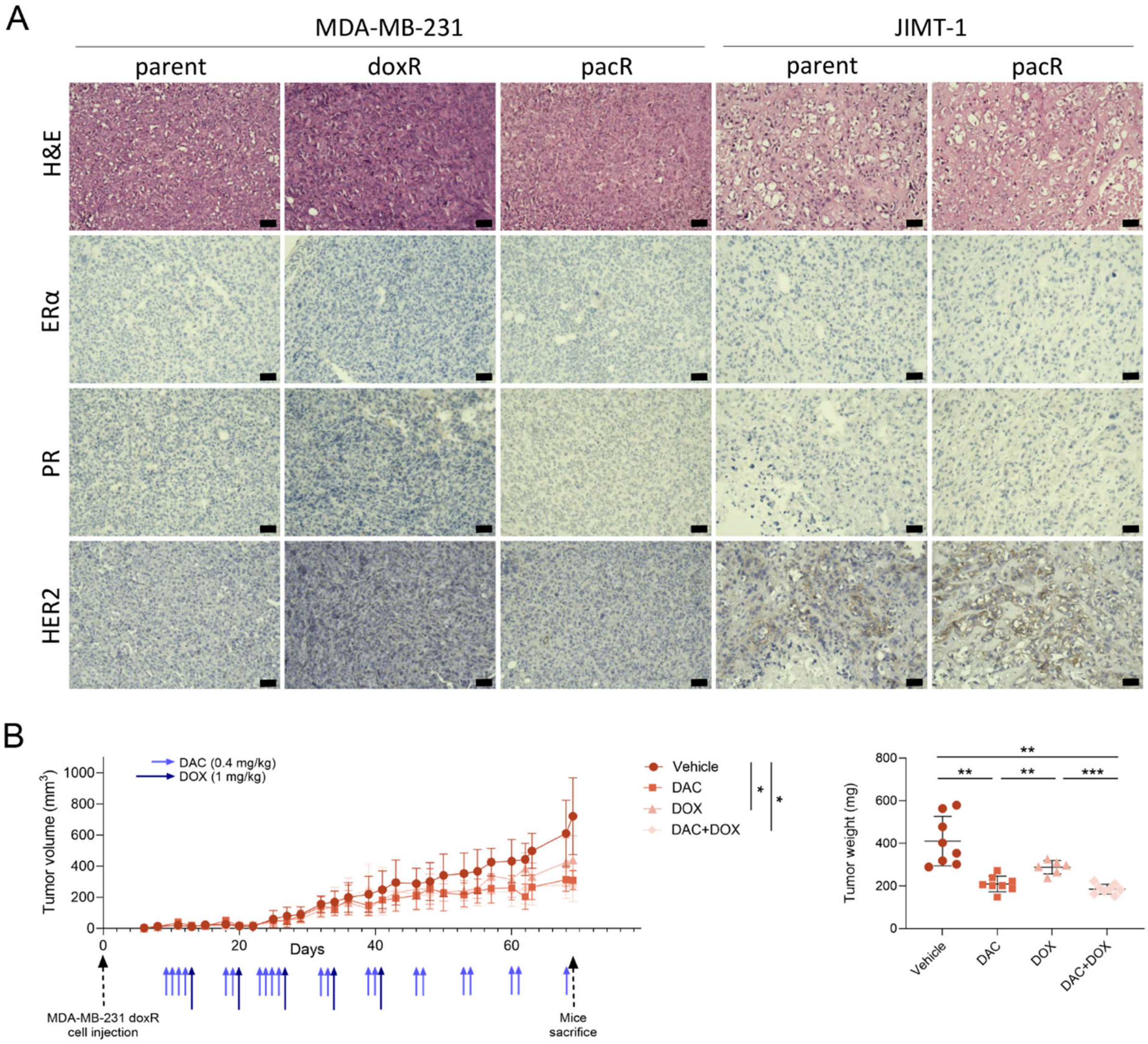


**Fig. S3.** Expression of hormone receptors and HER2 in cell line-derived xenografts, and evaluation of monotherapy and combination therapy in a pilot *in vivo* doxorubicin-resistant (doxR) model. (A) Representative immunohistochemical images of ERα, PR, and HER2 captured at 200× magnification (50 µm scale bar). (B) Pilot experiment showing therapeutic efficacy of decitabine (DAC), doxorubicin (DOX), and their combination in MDA-MB-231 doxR xenograft model following the scheme with prolonged treatment indicated by arrows: DAC (light blue), DOX (dark blue). DOX treatment was discontinued after five doses due to the cumulative dose limit being reached. Abbreviations: H&E, hematoxylin and eosin, ERα, estrogen receptor α, PR, progesterone receptor, HER2, human epidermal growth factor receptor 2.
